# Molecular framework for TIR1/AFB-Aux/IAA-dependent auxin sensing controlling adventitious rooting in Arabidopsis

**DOI:** 10.1101/518357

**Authors:** Abdellah Lakehal, Salma Chaabouni, Emilie Cavel, Rozenn Le Hir, Alok Ranjan, Zahra Raneshan, Ondřej Novák, Daniel I. Păcurar, Irene Perrone, François Jobert, Laurent Gutierrez, Laszlo Bakò, Catherine Bellini

## Abstract

In *Arabidopsis thaliana*, canonical auxin-dependent gene regulation is mediated by 23 transcription factors from the AUXIN RESPONSE FACTOR (ARF) family interacting with 29 auxin/indole acetic acid repressors (Aux/IAA), themselves forming coreceptor complexes with one of six TRANSPORT INHIBITOR1/AUXIN-SIGNALLING F-BOX (TIR1/AFB) PROTEINS. Different combinations of co-receptors drive specific sensing outputs, allowing auxin to control a myriad of processes. Considerable efforts have been made to discern the specificity of auxin action. However, owing to a lack of obvious phenotype in single loss-of-function mutants in *Aux/IAA* genes, most genetic studies have relied on gain-of-function mutants, which are highly pleiotropic. Using loss-of-function mutants, we show that three Aux/IAA proteins interact with ARF6 and/or ARF8, which we have previously shown to be positive regulators of AR formation upstream of jasmonate, and likely repress their activity. We also demonstrate that *TIR1* and *AFB2* are positive regulators of adventitious root formation and suggest a dual role for TIR1 in the control of JA biosynthesis and conjugation, as revealed by upregulation of several JA biosynthesis genes in the *tir1-1* mutant. We propose that in the presence of auxin, TIR1 and AFB2 form specific sensing complexes with IAA6, IAA9 and/or IAA17 that modulate JA homeostasis to control AR initiation.

## INTRODUCTION

In *Arabidopsis thaliana*, auxin-dependent gene regulation is mediated by the 23 members of the AUXIN RESPONSE FACTOR (ARF) family of transcription factors, which can either activate or repress transcription (Chapman and Estelle, 2009; Guilfoyle and Hagen, 2007). Interaction studies have shown that most of the 29 auxin/indole-3-acetic acid (Aux/IAA) inducible proteins can interact with ARF activators (Guilfoyle and Hagen, 2007; Vernoux et al., 2011). Aux/IAAs mediate recruitment of the TOPLESS corepressor (Szemenyei et al., 2008) and act as repressors of transcription of auxin-responsive genes. When the auxin level rises, it triggers interaction of the two components of the auxin co-receptor complex, an F-box protein from the TRANSPORT INHIBITOR1/AUXIN-SIGNALLING F-BOX PROTEIN (TIR1/AFB) family and an Aux/IAA protein, promoting ubiquitination and 26S-mediated degradation of the latter. Degradation of the Aux/IAA protein releases the ARF activity and subsequent activation of the auxin response genes (Wang and Estelle, 2014; Weijers and Wagner, 2016). TIR1/AFBs show different affinities for the same Aux/IAA (Calderon Villalobos et al., 2012; Parry et al., 2009), suggesting that different combinations of TIR1/AFB receptors may partially account for the diversity of auxin response. In addition, it has been shown that most Aux/IAAs can interact with many Aux/IAAs and ARFs in a combinatorial manner, increasing the diversity of possible auxin signaling pathways that control many aspects of plant development and physiology (Boer et al., 2014; Guilfoyle and Hagen, 2012; Korasick et al., 2014; Nanao et al., 2014; Vernoux et al., 2011; Weijers et al., 2005). Several studies have suggested specialized functions for some of the ARF and IAA combinations during embryo development (Hamann et al., 2002), lateral root (LR) development (De Rybel et al., 2010; De Smet et al., 2010; Fukaki et al., 2002; Lavenus et al., 2013; Tatematsu et al., 2004), phototropism (Sun et al., 2013) and fruit development (Wang et al., 2005). However, most of these studies involved characterization of gain-of-function stabilizing mutations, which limited identification of more specialized functions for individual Aux/IAA genes. To date, genetic investigations of Aux/IAA genes have been hampered by the lack of obvious phenotype in the loss-of-function mutants (Overvoorde et al., 2005). Nevertheless, recent careful characterization of a few of the mutants identified more precise functions in primary or LR development for *IAA3* or *IAA8* (Arase et al., 2012; Dello Ioio et al., 2008) or in the response to environmental stresses for *IAA3*, *IAA5*, *IAA6* and *IAA19* (Orosa-Puente et al., 2018; Shani et al., 2017).

To decipher the role of auxin in the control of adventitious root (AR) development, which is a complex trait with high phenotypic plasticity (Bellini et al., 2014; Geiss et al., 2009), we previously identified a regulatory module composed of three *ARF* genes (two activators *AFR6* and *ARF8*, and one repressor *ARF17*) and their regulatory microRNAs (miR167 and miR160) (Gutierrez et al., 2009). These genes display overlapping expression domains, interact genetically and regulate each other’s expression at transcriptional and post-transcriptional levels by modulating the availability of their regulatory microRNAs miR160 and miR167 (Gutierrez et al., 2009). The three ARFs control the expression of three auxin inducible *Gretchen Hagen 3* (*GH3*) genes encoding acyl-acid-amido synthetases (GH3.3, GH3.5 and GH3.6) that inactivate jasmonic acid (JA), an inhibitor of AR initiation in Arabidopsis hypocotyls ((Gutierrez et al., 2012) and Supplemental Figure 1A). In a yeast two-hybrid system, ARF6 and ARF8 proteins were shown to interact with almost all Aux/IAA proteins (Vernoux et al., 2011). Therefore, we propose a model in which increased auxin levels facilitate formation of a coreceptor complex with at least one TIR1/AFB protein and subsequent degradation of Aux/IAAs (Supplemental Figure 1B), thereby releasing the activity of ARF6 and ARF8 and the transcription of *GH3* genes. In the present work, we describe identification of members of the potential co-receptor complexes involved in this pathway. Using loss-of-function mutants, we demonstrate that *TIR1* and *AFB2* are positive regulators, whereas *IAA6*, *IAA9* and *IAA17* are negative regulators of AR formation. We suggest that TIR1 and AFB2 form co-receptor complexes with at least three Aux/IAA proteins (IAA6, IAA9 and IAA17), which negatively control *GH3.3*, *GH3.5* and *GH3.6* expression by repressing the transcriptional activity of ARF6 and ARF8, thereby modulating JA homeostasis and consequent AR initiation. In addition, we show that several genes involved in JA biosynthesis are upregulated in the *tir1-1* mutant, suggesting a probable dual role of TIR1 in both the biosynthesis and conjugation of jasmonate.

## RESULTS

### TIR1 and AFB2 but not other AFB proteins control adventitious root initiation in Arabidopsis hypocotyls

To assess the potential contributions of different TIR/AFB proteins to regulation of adventitious rooting in Arabidopsis, we analyzed AR formation in *tir1-1*, *afb1-3*, *afb2-3*, *afb3-4*, *afb4-8*, *afb5-5* single knockout (KO) mutants and double mutants using previously described conditions ((Gutierrez et al., 2009; Sorin et al., 2005) and Figure 1A). The average number of ARs developed by *afb1-3*, *afb3-4*, *afb4-8, afb5-5* single mutants and *afb4-8afb5-5* double mutants did not differ significantly from the average number developed by wild-type seedlings (Figure 1A). These results suggest that AFB1, AFB3, AFB4 and AFB5 do not play a significant role in AR initiation. In contrast, *tir1-1* and *afb2-3* single mutants produced 50% fewer ARs than the wild-type plants and the *tir1-1afb2-3* double mutant produced even fewer, indicating an additive effect of the mutations (Figure 1A). The *afb1-3afb2-3* and *afb2-3afb3-4* double mutants retained the same phenotype as the *afb2-3* single mutant, confirming a minor role, if any, of AFB1 and AFB3 in AR initiation. We also checked the root phenotype of the *tir1-1* and *afb2-3* single mutants and *tir1-1afb2-3* double mutant under the growth conditions used. No significant differences were observed in the primary root length (Supplemental Figure 1A), but the number of LRs was slightly but significantly decreased in both the *tir1-1* and *afb2-3* single mutants and dramatically decreased in the double mutant (Supplemental Figure 1B), as already shown by others (Dharmasiri et al., 2005b; Parry et al., 2009). This resulted in a reduction of the LR density in all genotypes (Supplemental Figure 1C), confirming the additive and pleiotropic role of the TIR1 and AFB2 proteins.

**Figure 1:**
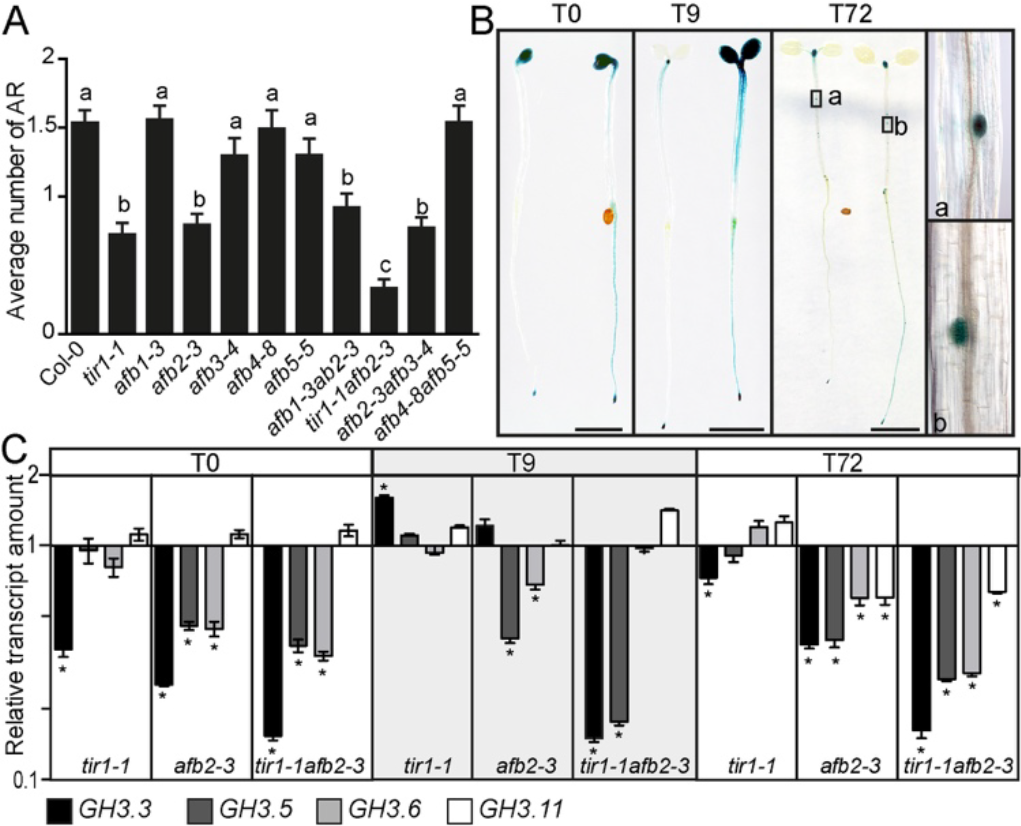
TIR1 and AFB2 control adventitious root initiation by modulating *GH3.3*, *GH3.5* and *GH3.6* expression. (A) Average numbers of adventitious roots in *tir/afb* mutants. Seedlings were first etiolated in the dark until their hypocotyls were 6 mm long and then transferred to the light for 7 days. Data were obtained from 3 biological replicates; for each, data for at least 30 seedlings were pooled and averaged. Errors bars indicate ± SE. One-way ANOVA combined with Tukey’s multiple comparison post-test indicated that only mutations in the *TIR1* and *AFB2* genes significantly affected the initiation of adventitious roots (*n*>30; *P* < 0.001). (B) Expression pattern of TIR1 and AFB2 proteins. GUS staining of *tir1-1pTIR1:cTIR1-GUS* and *afb2-3AFB2:cAFB2-GUS* translational fusions (arranged from left to right in each panel) in seedlings grown in the dark until their hypocotyls were 6 mm long (T0) and 9 h (T9) and 72 h (T72) after their transfer to the light. (a) and (b) Close-ups from hypocotyl regions shown for T72. (C) Quantification by qRT-PCR of *GH3.3*, *GH3.5* and *GH3.6* transcripts in hypocotyls of *tir1-1* and *afb2-3* single mutants and the *tir1-1afb2-3* double mutant. mRNAs were extracted from hypocotyls of seedlings grown in the dark until the hypocotyl reached 6 mm (T0) and after their transfer to the light for 9 h or 72 h. The gene expression values are relative to the expression in the wild type, for which the value was set to 1. Error bars indicate ± SE obtained from three independent biological replicates. One-way ANOVA combined with Dunnett’s multiple comparison test indicated that in some cases, the relative amount of mRNA was significantly different from the wild type (denoted by *, *P* < 0.001; *n* = 3).

### TIR1 and AFB2 proteins are expressed in young seedlings during AR initiation

To analyze the expression pattern of the TIR1 and AFB2 proteins during the early stages of AR initiation and development, plants expressing the translational fusions *pTIR:cTIR1:GUS* or *pAFB2:cAFB2:GUS* were grown as previously described (Gutierrez et al., 2009). At time 0 (T0), i.e., in etiolated seedlings just before transfer to the light, the TIR1:GUS and AFB2:GUS proteins were strongly expressed in the root apical meristem, apical hook and cotyledons. Interestingly AFB2:GUS was also detected in the vascular system of the root and the hypocotyl, whereas TIR1:GUS was not detectable in those organs (Figure 1B). Nine hours after transfer to the light, TIR1:GUS protein disappeared from the cotyledons but was still strongly expressed in the shoot and root meristems. Its expression was increased slightly in the upper part of the hypocotyl. In contrast, AFB2:GUS was still highly detectable in the shoot and root meristems, cotyledons and vascular system of the root. In addition, its expression was induced throughout almost the entire hypocotyl (Figure 1B). Seventy-two hours after transfer to the light, TIR1:GUS and AFB2:GUS showed almost the same expression pattern, which was reminiscent of that previously described in light grown seedlings (Parry et al., 2009). None of the proteins were detectable in the cotyledons. However, they were present in the shoot meristem and young leaves and the apical root meristem. In the hypocotyl and root, the TIR1:GUS and AFB2:GUS proteins were mainly detectable in the AR and LR primordia (Figure 1B).

### TIR1 likely controls both JA biosynthesis and conjugation, whereas AFB2 preferentially controls JA conjugation during adventitious root initiation

Based on our model (Supplemental Figure 1A and B), one would expect to see downregulation of the *GH3.3*, *GH3.5* and *GH3.6* genes in the *tir1-1*, *afb2-3* single mutants and *tir1-1afb2-3* double mutant. Therefore, we analyzed the relative transcript amount of the three *GH3* genes in these mutants (Figure 1C). *GH3-11/JAR1*, which conjugates JA into its bioactive form jasmonoyl-L-isoleucine (JA-Ile), was used as a control. Its expression was only slightly downregulated in the *afb2-3* single mutant and *tir1-1afb2-3* double mutant at T72 (Figure 1C), whereas expression of the other three *GH3* genes was significantly reduced in the *afb2-3* single mutant and *tir1-1afb2-3* double mutant at all timepoints (Figure 1C). In the *tir1-1* single mutant, only *GH3.3* was significantly downregulated at T0 and slightly downregulated at T72 (Figure 1C), but an additive effect of the *tir1-1* mutation on the expression *GH3.3, GH3.5* and *GH3.6* was observed in the *tir1-1afb2-3* double mutant at all timepoints (Figure 1C), suggesting a redundant role for TIR1 in the regulation of JA conjugation. Our results suggest that AFB2 likely controls AR initiation by regulating JA homeostasis through the *ARF6/ARF8* auxin signaling module (as shown in Supplemental Figure 1) and that TIR1, besides its redundant function in JA conjugation, might have another role in controlling ARI by regulating other hormone biosynthesis and/or signaling cascades. To test this hypothesis, we quantified endogenous free salicylic acid (SA), free IAA, free JA and JA-Ile (Figure 2A to D) in the hypocotyls of wild-type seedlings and seedlings of the *tir1-1*, *afb2-3* single mutants and *tir1-1afb2-3* double mutant. No significant differences in SA content were observed between the wild type and mutants (Figure 2A). A slight but significant increase in free IAA content was observed at T0 in all three mutants compared to the wild type (Figure 2B), but only in the *tir1-1afb2-3* double mutant at 9 and 72 hours after transfer to the light (Figure 2B). This slight increase in the free IAA content can be explained by feedback regulation as a consequence of downregulation of the auxin signaling pathway in the mutants. At T0 and T9, a significant increase in free JA was observed in both the *tir1-1* and *afb2-3* single mutants compared to the wild type but not in the double mutant *tir1-1afb2-3* (Figure 2C). The bioactive form JA-Ile was significantly accumulated in the single mutants at all three time points but accumulated only at T9 in the double mutant *tir1-1afb2-3* (Figure 2D). The fact that JA and JA-Ile did not accumulate in the double mutant can be explained by negative feedback loop regulation of JA homeostasis. Accumulation of JA and JA-Ile in the *afb2-3* mutant was expected since the three GH3 conjugating enzymes were found to be downregulated (Figure 1C), but we did not *a priori* expect the same level of accumulation for the *tir1-1* mutant. These results prompted us to check the expression of JA biosynthesis genes in the mutants to investigate the potential role of TIR1 and/or AFB2 in the control of JA biosynthesis. The relative transcript amounts of seven key genes involved in JA biosynthesis were analyzed by qRT-PCR in the hypocotyls of wild-type, *tir1-1*, *afb2-3* and *tir1-1afb2-3* seedlings grown under adventitious rooting conditions (Figure 2E to G). In etiolated seedlings (T0), *OPCL1*, *OPR3*, *AOC2* were significantly upregulated in the *tir1-1* mutant compared to the wild type, whereas *LOX2* was downregulated. In the *afb2-3* mutant, no significant differences were observed except for *LOX2* and *AOC1*, which were downregulated compared to the wild type. In the double mutant, *LOX2* and *AOC2* were significantly upregulated (Figure 2E). Nine hours after transfer to the light (T9), five (*OPCL1*, *OPR3*, *LOX2*, *AOC2, AOC3*) out of the seven biosynthesis genes were significantly upregulated in the single *tir1-1* mutant and four of them (*OPCL1*, *OPR3*, *LOX2*, *AOC2*) were upregulated in the *tir1-1afb2-3* double mutant (Figure 2F). Only *AOC3* and *AOC4* were upregulated in the *afb2-3* mutant at T9 (Figure 2F). At T72, only *LOX2* was significantly upregulated in all three mutants (Figure 2G). In conclusion, expression of JA biosynthesis genes was more significantly upregulated in the single *tir1-1* mutant than in the *afb2-3* mutant during AR initiation. Therefore, we propose that TIR1 and AFB2 control JA homeostasis, with a major role for TIR1 in the control of JA biosynthesis and a major role for AFB2 in the control of JA conjugation through the *ARF6/ARF8* auxin signaling module.

**Figure 2:**
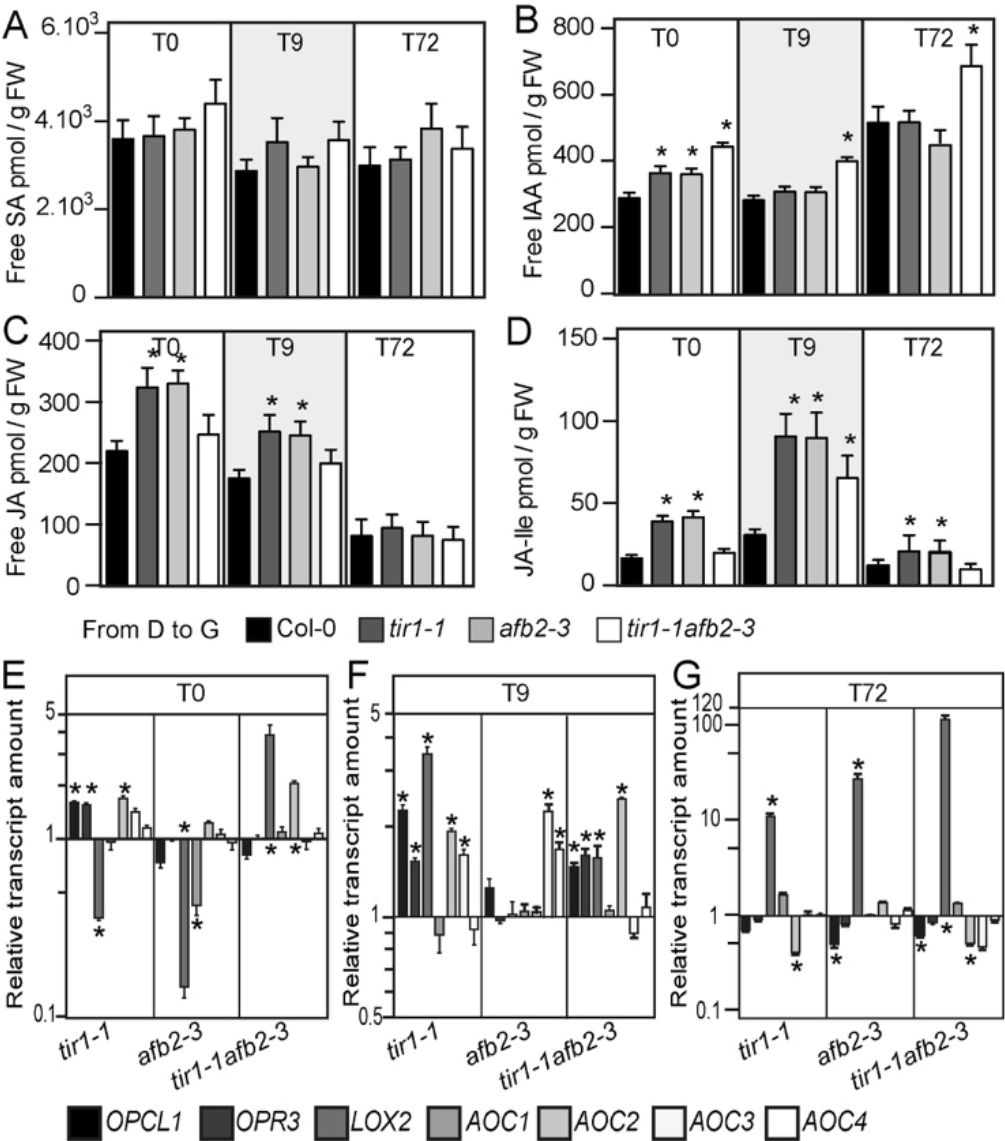
TIR1 and AFB2 control adventitious root initiation by modulating jasmonate homeostasis. (A) to (D) The endogenous contents of free IAA (D), free SA (B), free JA (C) and JA-Ile (D) were quantified in the hypocotyls of wild type Col-0, single mutants *tir1-1* and *afb2-3* and double mutant *tir1-1afb2-3* seedlings grown in the dark until the hypocotyl reached 6 mm (T0) and after their transfer to the light for 9 h (T9) or 72 h (T72). Error bars indicate ± SD of six biological replicates. One-way ANOVA combined with Dunnett’s multiple comparison test indicated that in some cases, values were significantly different from those of the wild-type Col-0 (denoted by *, *P* < 0.05; *n* = 6). (E) to (G) Relative transcript amount of genes involved in JA biosynthesis (*OPCL1, OPR3*, *LOX2*, *AOC1, AOC2, AOC3, AOC4*). The transcript amount was assessed by qRT-PCR using mRNAs extracted from hypocotyls of seedlings grown in the dark until the hypocotyl reached 6 mm (T0) and after their transfer to the light for 9 h (T9) or 72 h (T72). The gene expression values are relative to the expression in the wild type, for which the value was set to 1. Error bars indicate ± SE obtained from three independent biological replicates. One-way ANOVA combined with the Dunnett’s multiple comparison test indicated that in some cases, the relative amount of mRNA was significantly different from the wild type (denoted by *, *P* < 0.001; *n* = 3).

### A subset of Aux/IAA proteins regulate adventitious root initiation in Arabidopsis hypocotyls

ARF6 and ARF8 are two positive regulators of AR initiation (Gutierrez et al., 2009; Gutierrez et al., 2012) and their transcriptional activity is known to be regulated by Aux/IAA genes. To gain further insight into the auxin sensing machinery and complete our proposed signaling module involved in AR initiation, we attempted to identify potential Aux/IAA proteins that interact with ARF6 and/or ARF8. In 2011, Vernoux *et al.* (2011) conducted a large-scale analysis of the Aux/IAA-ARF network using a high-throughput yeast two-hybrid approach. They showed that ARF6 and ARF8 belong to a cluster of proteins that can interact with 22 of the 29 Aux/IAA genes (Vernoux et al., 2011). However, this does not help much to restrict the number of genes of interest. Hence, to elucidate which Aux/IAAs can interact with ARF6 and ARF8 during AR formation, we looked at those most expressed in the hypocotyl and assessed the expression of the 29 *Aux/IAA* genes in different organs (cotyledons, hypocotyl and roots) of 7-day-old light-grown seedlings using qRT-PCR (Supplemental Figure 3). With the exception of *IAA15*, we detected a transcript for all *IAA* genes in all organs tested (Supplemental Figure 3). We observed that 18 *IAA* genes were more expressed in the hypocotyl compared to cotyledons or roots (*IAA1, IAA2, IAA3, IAA4, IAA5, IAA6, IAA7, IAA8, IAA9, IAA10, IAA13, IAA14, IAA16, IAA19, IAA26, IAA27, IAA30, IAA31*), 4 *IAA* genes were more expressed in the hypocotyl and the root (*IAA17*, *IAA20, IAA28, IAA33*) and 6 genes were more expressed in the cotyledons (*IAA11, IAA12, IAA18, IAA29, IAA32, IAA34*). To assess the potential contributions of different *IAA* genes in the regulation of AR, we obtained KO mutants available for nine of the *Aux/IAA* genes that displayed high expression in the hypocotyl (*iaa3/shy2-24, iaa4-1, iaa5-1, iaa6-1, iaa7-1, iaa8-1, iaa9-1, iaa14-1, iaa30-1*), two of the genes which had high expression in both the hypocotyl and root (*iaa17-6*, *iaa28-1, iaa33-1*) and we added two KO mutants with genes whose expression was lower in the hypocotyl and root (*iaa12-1* and *iaa29-1*).

We analyzed AR formation in the *iaa* KO mutants under previously described conditions (Gutierrez et al., 2009; Sorin et al., 2005). Interestingly, six mutants (*iaa5-1*, *iaa6-1*, *iaa7-1*, *iaa8-1*, *iaa9-1* and *iaa17-6*) produced significantly more ARs than the wild type, whereas all the other mutants did not show any significant difference compared to the wild type (Figure 3A). The primary root length and LR number were not affected in mutants *iaa5-1*, *iaa6-1* and *iaa8-1* (Supplemental Figure 2D to F), whereas *iaa9-1* and *iaa17-6* showed a slightly shorter primary root and fewer LRs than the wild type (Supplemental Figure 2D and E) but the LR density was not affected (Supplemental Figure 2F). In contrast, *iaa7-1* had a slightly but significantly longer primary root as well as fewer LRs, which led to a slightly but significantly decreased LR density (Supplemental Figure 2F). These results strongly suggest that *IAA5*, *IAA6*, *IAA7*, *IAA8*, *IAA9* and *IAA17* are involved in the control of AR formation and substantiate our hypothesis that only a subset of *Aux/IAA* genes regulate the process of AR formation.

**Figure 3:**
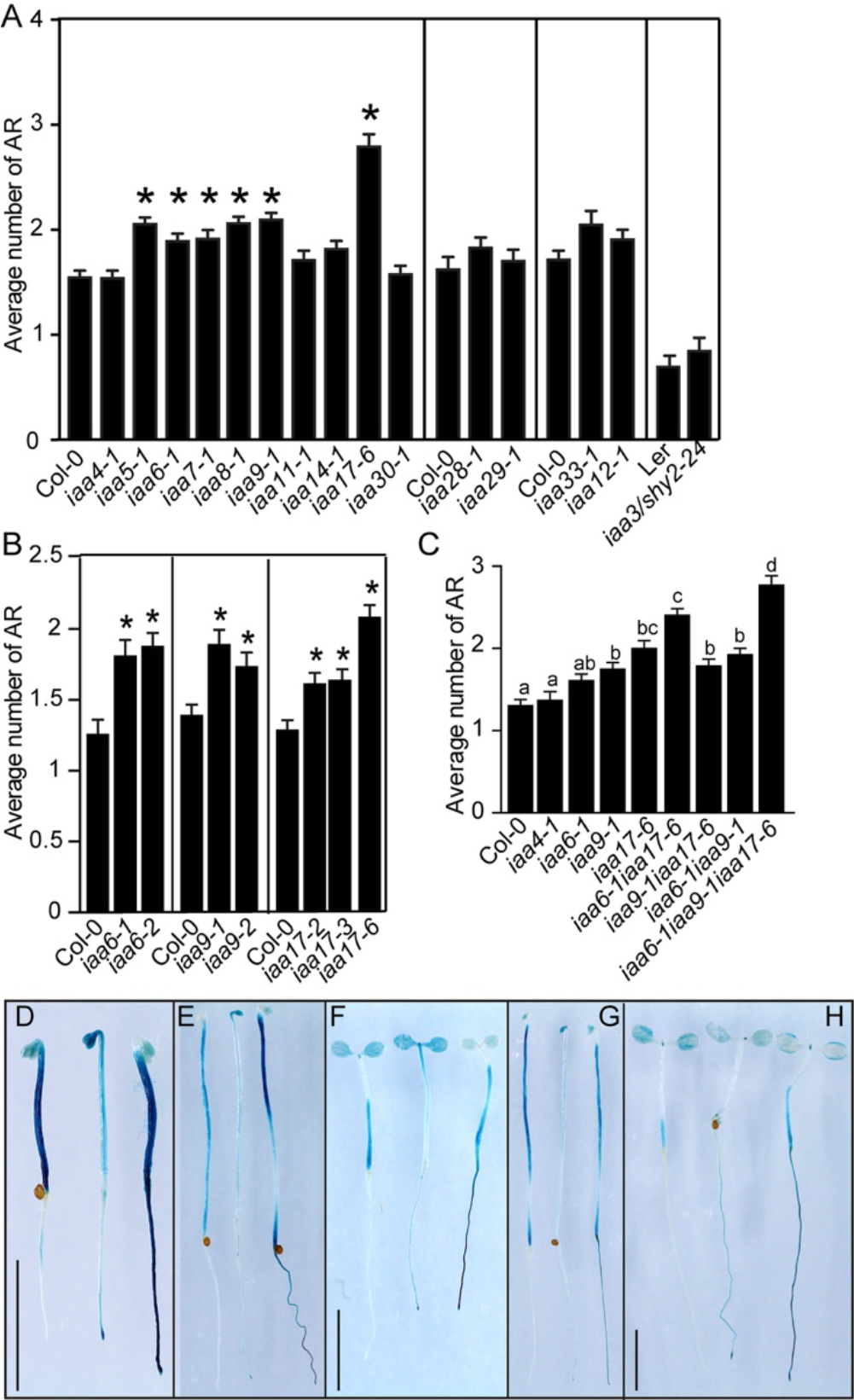
*IAA6*, *IAA9* and *IAA17* are involved in the control of adventitious root initiation. (A) Average numbers of ARs assessed in 15 *aux/iaa* knockout mutants. (B) Average numbers of ARs in *iaa6-1, iaa6-2*, *iaa9-1, iaa9-2*, *iaa17-2, iaa17-3* and *iaa17-6* mutant alleles. (C) Average numbers of ARs in single *iaa6-1*, *iaa9-1* and *iaa17-6* single, double and triple mutants. (A) to (C) Seedlings were first etiolated in the dark until their hypocotyls were 6 mm long and then transferred to the light for 7 days. Data were obtained from 3 biological replicates; for each, data for at least 30 seedlings were pooled and averaged. Errors bars indicate ± SE. In (A) and (B), one-way ANOVA combined with Dunnett’s multiple comparison post-test indicated that in some cases, differences observed between the mutants and the corresponding wild type were significant (denoted by *, *P* < 0.001, *n* > 30). In (C), one-way ANOVA combined with Tukey’s multiple comparison post-test indicated significant differences (denoted by different letters, *P* < 0.001, *n* > 30) (D) to (H) Expression pattern of *IAA6*, *IAA9* and *IAA17* during the initial steps of AR formation. GUS staining of *promIAA6:GUS*, *promIAA9:GUS* and *promIAA17:GUS* (arranged from left to right in each panel) in seedlings grown in the dark until their hypocotyls were 6 mm long (D), after additional 48 h (E) and 72 h (G) after in the dark, and 48 h (F) and 72 h (H) after their transfer to the light. Bars = 5 mm.

### IAA6, IAA9 and IAA17 proteins interact with ARF6 and ARF8 proteins

To establish whether these targeted proteins were effective partners of ARF6 and ARF8, we performed co-immunoprecipitation (CoIP) in protoplasts transfection assays. Arabidopsis protoplasts were transfected with plasmids expressing cMyc- or HA-tagged AuxIAA and ARF proteins according to the protocol described in the Materials and Methods (Magyar et al., 2005). The presence of the putative ARF/AuxIAA complex was tested by western blotting with anti-HA or anti-c-Myc antibodies and only interactions with *IAA6*, *IAA9* and *IAA17* were detected (Figure 5A to E): IAA6 and IAA17 interacted with ARF6 and ARF8 (Fig. 5*A*, *B*, *D* and *E*), whereas IAA9 interacted only with ARF8 (Figure 5C). These results were confirmed by a bimolecular fluorescence complementation (BiFC) assay (Figure 5I to M)

### ARF6 but not ARF8 can form a homodimer

Recent interaction and crystallization studies have shown that ARF proteins dimerize *via* their DNA-binding domain (Boer et al., 2014) and interact not only with Aux/IAA proteins but potentially also with themselves or other ARFs *via* their PB1 domain with a certain specificity (Vernoux et al., 2011). Therefore, we also used CoIP and BiFC assays and tagged versions of the ARF6 and ARF8 proteins to check whether they could form homodimers and/or a heterodimer. Our results (Figure 5G, H, O and P) agreed with a previously published yeast two-hybrid interaction study (Vernoux et al., 2011), which showed that ARF6 and ARF8 do not interact to form a heterodimer and that ARF8 does not homodimerize. In contrast, we showed that ARF6 protein can form a homodimer (Figure 5F and N), suggesting that ARF6 and ARF8, although redundant in controlling the expression of *GH3.3, GH3.5* and *GH3.6* genes (30), might have a specificity of action.

### *IAA6*, *IAA9* and *IAA17* act redundantly to control adventitious root initiation

Because we found an interaction only with the IAA6, IAA9 and IAA17 proteins, we continued to characterize the role of their corresponding genes. All three single *iaa* mutants showed a significant and reproducible AR phenotype. Nevertheless, because extensive functional redundancy has been shown among *Aux/IAA* gene family members (Overvoorde et al., 2005), it was important to confirm the phenotype in at least a second allele (Figure 3B). We also generated the double mutants *iaa6-1iaa9-1*, *iaa6-1iaa17-6* and *iaa9-1iaa17*-6 and the triple mutant *iaa6-1iaa9-1iaa17-6* and analyzed their phenotype during AR formation (Figure 3C). Mutant *iaa4-1* was used as a control showing no AR phenotype. Except for the *iaa6iaa17-6* double mutant, which showed an increased number of AR compared to the single mutants, the other two double mutants were not significantly different from the single mutants (Figure 3C). Nevertheless, we observed a significant increase of the AR number in the triple mutants compared to the double mutants, suggesting that these genes act redundantly in the control of AR initiation (Figure 3C) but do not seem to be involved in the control of the PR or LR root growth as shown on (Supplemental Figure 2G-I). We also characterized the expression of *IAA6*, *IAA9* and *IAA17* during the early steps of AR formation using transcriptional fusion constructs containing a ß-glucuronidase (GUS) coding sequence fused to the respective promoters. At time T0 (i.e., etiolated seedlings prior to transfer to the light) (Figure 3D), *promIAA6:GUS* was strongly expressed in the hypocotyl, slightly less expressed in the cotyledons and only weakly expressed in the root; *promIAA9:GUS* was strongly expressed in the cotyledons, hook and root tips and slightly less in the hypocotyl and root; *promIAA17:GUS* was strongly expressed in the hypocotyl and root, slightly less in the cotyledons and, interestingly, was excluded from the apical hook (Figure 3D). Forty-eight and seventy-two hours after transfer to the light, a decrease in GUS staining was observed for all the lines (Figure 3F and H), but only for *IAA9* when the seedlings were kept longer in the dark (Figure 3E and G). These results suggest that light negatively regulates the expression of *IAA6* and *IAA17* while the expression of IAA9 seem to depend on the developmental stage.

### *IAA6, IAA9* and *IAA17* negatively control expression of *GH3.3*, *GH3.5* and *GH3.6*

In our model, auxin stimulates adventitious rooting by inducing *GH3.3*, *GH3.5* and *GH3.6* gene expression *via* the positive regulators ARF6 and ARF8 (Supplemental Figure 1). Although we confirmed an interaction between IAA6, IAA9 and IAA17 with ARF6 and/or ARF8, it was important to demonstrate whether disrupting the expression of one of those genes would result in upregulation of *GH3* gene expression. Therefore, we performed qRT-PCR analysis of the relative transcript amounts of the three genes *GH3.3*, *GH3.5*, *GH3.6* in the hypocotyls of single mutants *iaa6-1*, *iaa9-1*, *iaa17-6* first etiolated and then transferred to the light for 72 h. The mutant *iaa4.1*, which had no phenotype affecting AR initiation (Figure 3A), was used as a control. Expression of *GH3.3*, *GH3.5* and *GH3.6* was upregulated in the *iaa9-1* mutant (Figure 4A), whereas only *GH3.3*, *GH3.5* were significantly upregulated in the *iaa6-1* and *iaa17-6* mutant (Figure 4A). In contrast, expression of *GH3.3*, *GH3.5* and *GH3.6* remained unchanged in the *iaa4-1* mutant (Figure 4A). These results confirm that IAA6, IAA9 and IAA17 are involved in the regulation of adventitious rooting through the modulation of *GH3.3*, *GH3.5* and *GH3.6* expression. To establish whether the *iaa6-1*, *iaa9-1* and *iaa17-6* mutations affected other *GH3* genes, the relative transcript amount of *GH3-10* and *GH3-11* was quantified. Notably, accumulation of *GH3.10* and *GH3.11/JAR1* transcripts was not significantly altered in the *iaa6-1*, *iaa9-1* and *iaa17-6* mutants but *GH3.10* was upregulated in the *iaa4-1 mutant* (Figure 4A). We concluded that *IAA6*, *IAA9* and *IAA17* negatively regulate *GH3.3*, *GH3.5* and *GH3.6* expression in the Arabidopsis hypocotyl during AR initiation.

**Figure 4:**
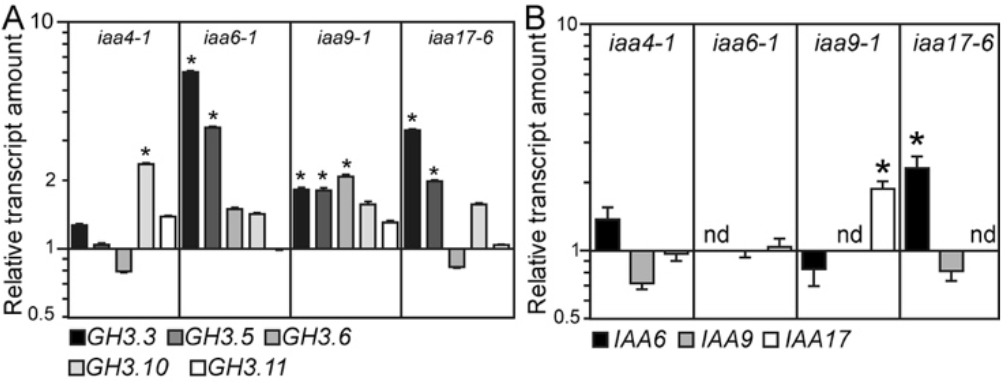
*IAA6*, *IAA9* and *IAA17* are involved in the control of adventitious root initiation upstream of *GH3.3*, *GH3.5* and *GH3.6*. (A) Relative transcript amount of *GH3.3*, *GH3.5*, *GH3.6*, *GH3.10* and *GH3.11* genes in hypocotyls of *iaa4-1, iaa6-1, iaa9-1* and *iaa17-6* single mutants. Relative transcript amount of *IAA6*, *IAA9* and *IAA17* genes in hypocotyls of *iaa4-1, iaa6-1, iaa9-1* and *iaa17-6* single mutants. In (A) and (B), mRNAs were extracted from hypocotyls of seedlings grown in the dark until the hypocotyl reached 6 mm and then transferred to the light for 72 h. Gene expression values are relative to expression in the wild type, for which the value was set to 1. Error bars indicate ± SE obtained from three independent biological replicates. One-way ANOVA combined with Dunnett’s multiple comparison test indicated that in some cases, the relative amount of mRNA was significantly different from the wild type (denoted by *, *P* < 0.001; *n* = 3).

**Figure 5:**
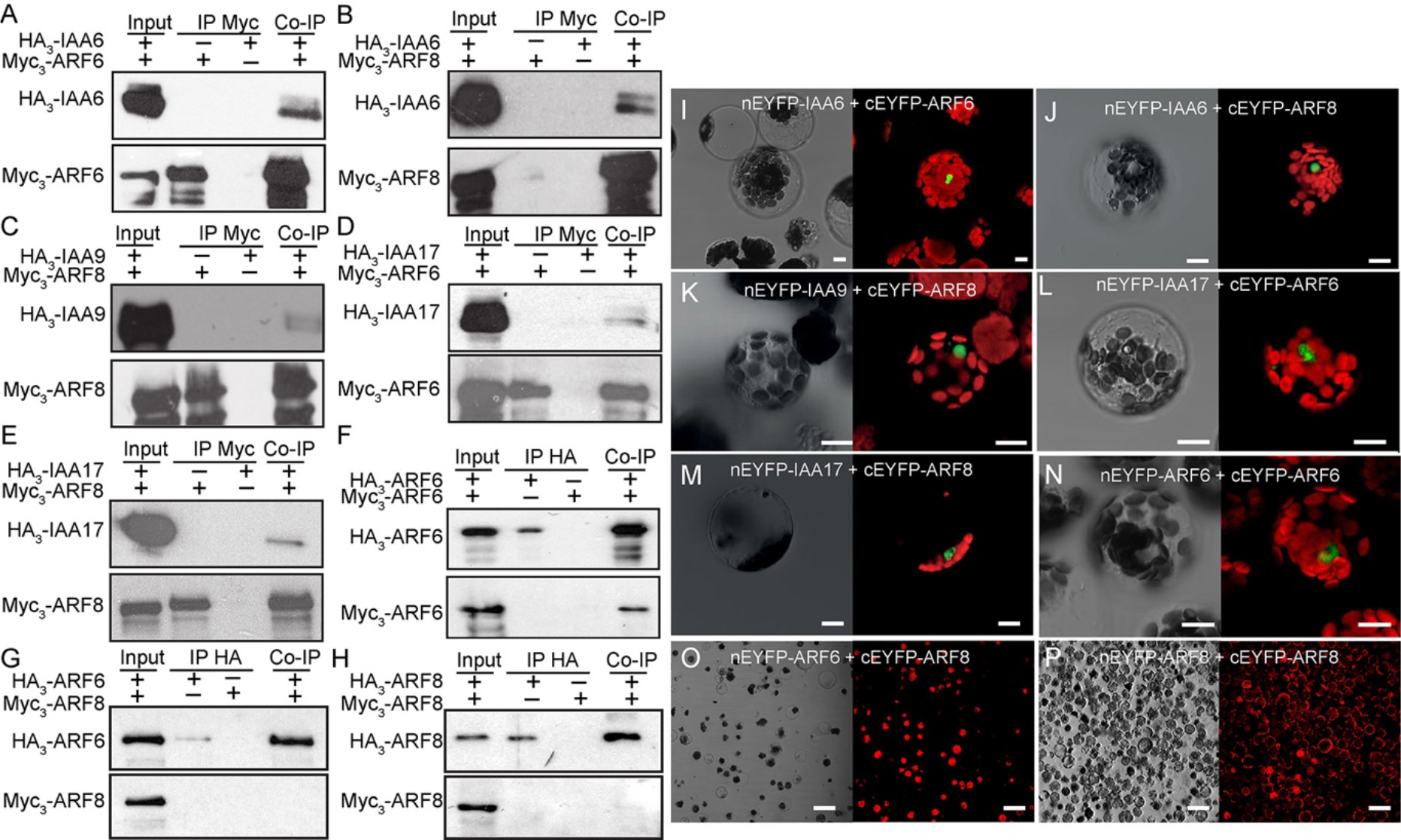
IAA6, IAA9 and IAA17 repressor proteins physically interact with ARF6 and/or ARF8, while ARF6 interacts with itself to form a homodimer. (A) to (E) Co-immunoprecipitation (CoIP) assay. Arabidopsis protoplasts were transfected with a HA_3_-tagged version of *IAA6*, *IAA9* or *IAA17* constructs and/or a c-Myc3-tagged version of *ARF6* or *ARF8* constructs. Proteins were immunoprecipitated with anti-Myc antibodies and submitted to anti-cMyc protein (lower panel) to confirm the presence of the ARF protein and to anti-HA gel-blot analysis to reveal the IAA partner (top panel). HA_3_-IAA6-cMyc-ARF6 (A), HA_3_-IAA6-cMyc-ARF8 (B), HA_3_-IAA9-cMyc-ARF8 (C), HA_3_-IAA17-cMyc-ARF6 (D), HA_3_-IAA17-cMyc-ARF6 (E). (F) to (H) Arabidopsis protoplasts were transfected with HA_3_-tagged and c-Myc3-tagged versions of *ARF6 and/or ARF8.* Proteins were immunoprecipitated with anti-HA antibodies and submitted to anti-HA protein (top panel) to confirm the presence of the ARF protein and to anti-cMyc antibody to reveal the ARF6 or ARF8 partner (top panel). Only ARF6 homodimer could be detected (F). (I) to (P) Confirmation of the interaction by bimolecular fluorescence complementation experiments (BiFC). Only Arabidopsis mesophyll protoplasts with intact plasma membranes, shown with bright-field light microscopy (left photo in each panel), tested positive for the presence of yellow fluorescence, indicating protein-protein interaction due to assembly of the split YFP, shown by confocal microscopy (right photo in each panel). (I) Cotransformation of 10 µg nEYFP-IAA6 and 10 µg ARF6-cEYFP into protoplasts generated yellow fluorescence (false-colored green) at the nucleus surrounded by chloroplast autofluorescence (false-colored red). Fluorescence was also observed after cotransformation of 10 µg of nEYFP-IAA6 and cEYFP-ARF8 (J); nEYFP-IAA9 and cEYFP-ARF8 (K); nEYFP-IAA17 and cEYFP-ARF6 (L); nEYFP-IAA17 and cEYFP-ARF8 (M), and nEYFP-ARF6 and cEYFP-ARF6 (N). No fluorescence was detected after cotransformation of 10 µg of nEYFP-ARF6 and cEYFP-ARF8 (O) or nEYFP-ARF8 and cEYFP-ARF8 (P). Bars = 10 µm.

We also checked a possible compensatory effect induced by the knockout of one the IAA genes. We performed qRT-PCR analysis of the relative transcript amounts of *IAA6*, *IAA9* and *IAA17* genes in the hypocotyl of each single mutant (Figure 4B). Interestingly, a mutation in the *IAA6* gene did not affect the expression of *IAA9* or *IAA17*, whereas *IAA17* was significantly upregulated in the hypocotyls of *iaa9-1* mutant seedlings. *IAA6* was upregulated in the hypocotyl of *iaa17-6* mutant seedlings and a mutation in *IAA4* did not affect the expression of any of the three *IAA* genes of interest (Figure 4B).

### *ARF6, ARF8* and *ARF17* are unstable proteins and their degradation is proteasome dependent

While transfecting Arabidopsis protoplasts for CoIP assays with open reading frames encoding individual cMyc- or HA-tagged versions of ARFs and Aux/IAAs, problems were encountered due to instability not only of the tagged Aux/IAA proteins but also of the tagged ARFs. It has previously been reported that like Aux/IAA proteins, ARFs may be rapidly degraded (Salmon et al., 2008). Therefore, we analyzed the degradation of HA_3_:ARF6, cMyc_3_:ARF8 and HA_3_:ARF17. We used HA_3_:ARF1, which was previously used as a control (Fig. 6A,E,F) (Salmon et al., 2008). Western blot analysis with protein extracts from transfected protoplasts using anti-HA or anti-cMyc antibodies showed that like ARF1, proteins ARF6, ARF8 and ARF17 were degraded. The HA_3_:ARF6 levels decreased dramatically within 30 minutes, indicating that ARF6 is a short-lived protein (Figure 6B), while the degradation rate of HA_3_:ARF17 was similar to that of HA_3_:ARF1 (Figure 6D) and cMyc_3_ARF8 appeared more stable (Figure 6C). To verify whether ARF6, ARF8 and ARF17 proteolysis requires activity of the proteasome for proper degradation, transfected protoplasts were incubated for 2 h in the presence or absence of 50 μM of a cell permeable proteasome-specific inhibitor, Z-Leu-Leu-Leu-CHO aldehyde (MG132), and the extracted proteins were analyzed by immunoblotting (Fig. 6E). The sample incubated with MG132 contained higher levels of HA_3_:ARF1, confirming the previously described proteasome-dependent degradation of ARF1 (34), and thereby the efficiency of the treatment. Similarly, HA_3_:ARF6, cMyc_3_ARF8 and HA_3_:ARF17 proteins accumulated in protoplasts treated with MG132, indicating that ARF6, ARF8 and ARF17 degradation is also proteasome dependent (Figure 6E). To further determine whether proteasome activity is necessary for ARF6, ARF8 and ARF17 protein degradation *in vivo*, one-week-old transgenic *in vitro* grown Arabidopsis seedlings expressing HA_3_:ARF1, cMyc_3_:ARF6, cMyc_3_:ARF8 and cMyc_3_:ARF17 were treated with MG132 or DMSO for 2 h prior to protein extraction. After western blotting, we observed that levels of HA_3_:ARF1, cMyc_3_:ARF6, cMyc_3_:ARF8 and cMyc_3_:ARF17 were enhanced by the addition MG132, confirming that their degradation is proteasome dependent in planta (Figure 6F).

**Figure 6:**
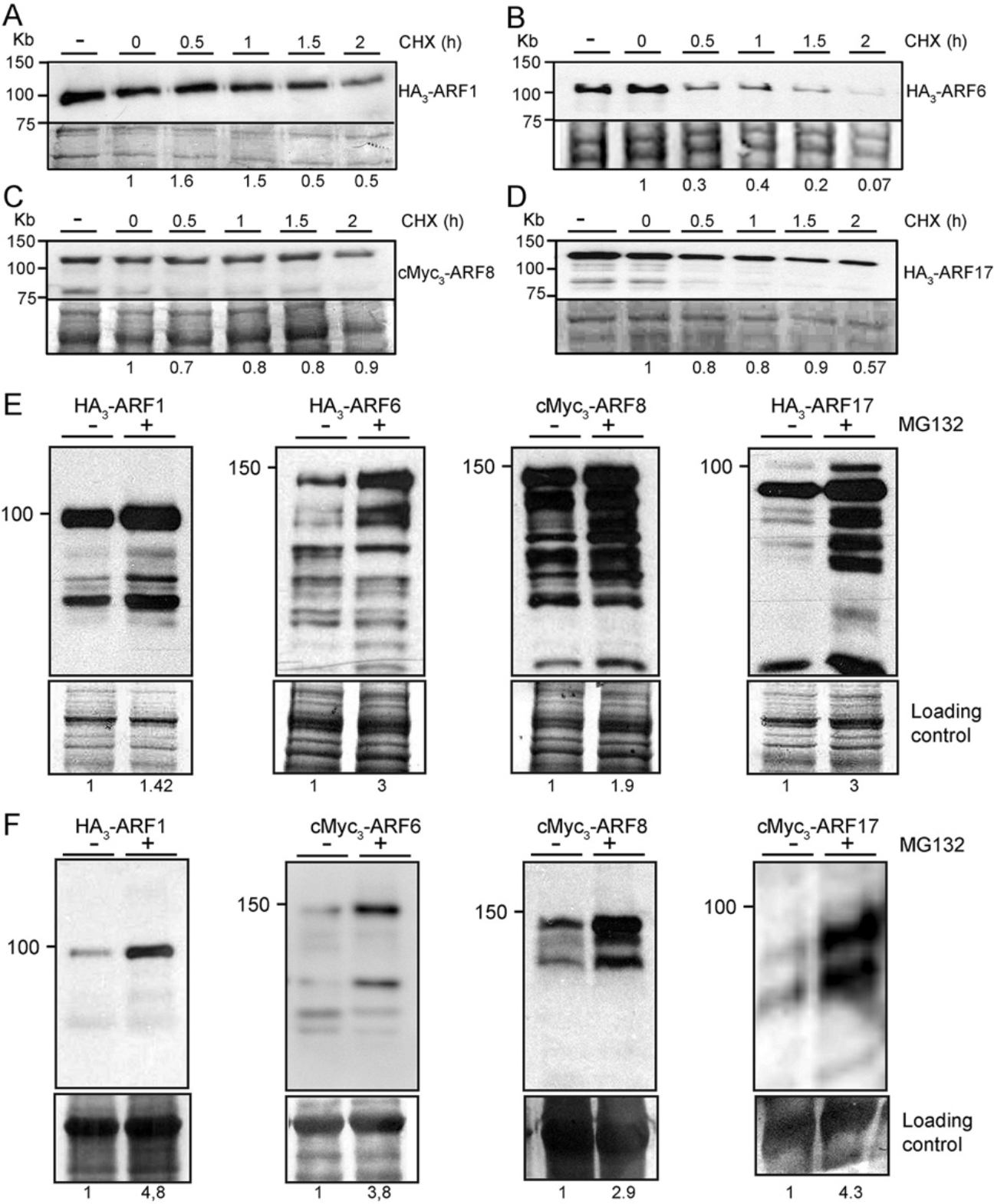
ARF6, ARF8 and ARF17 are unstable proteins whose degradation is proteasome dependent. (A) to (D) Degradation kinetics of ARF6, ARF8 and ARF17 proteins. Top panel: representative anti-HA or anti-c-Myc western blot performed on total protein from wild-type Col-0 protoplasts transformed with 5 μg of plasmid DNA expressing HA_3_- or cMyc3-tagged proteins and mock treated with DMSO (-) or treated with 200 μg/ml of cycloheximide. Lower panel: Amido Black staining of the membrane indicating protein loading. (E) Effect of MG132 on the degradation of the tagged ARF proteins in protoplasts. Top panel: representative anti-HA western blot performed on total protein from wild-type Col-0 protoplasts transformed with 5 μg of plasmid DNA expressing HA_3_- or cMyc3-ARF6, ARF8 and ARF17 or 15 μg of plasmid DNA expressing HA_3_-ARF1 treated with MG132 (+) or mock treated with DMSO (-) for 2 h. Lower panel: Amido Black staining of the membrane indicating protein loading. (F) Effect of MG132 on the degradation of the tagged ARF proteins *in Planta.* Top panel: representative western blot performed on total protein extracted from 7-day-old seedlings expressing HA_3_-ARF1, Myc_3_-ARF6, Myc_3_-ARF8 or Myc_3_-ARF17 treated with MG132 (+) or mock treated with DMSO (-) for 2 h. Lower panel: Amido Black staining of the membrane indicating protein loading. ImageJ (https://imagej.nih.gov/ij/) was used for densitometry imaging to analyze intensity of western blot bands. The ARFs staining intensities were quantified with the area of the major pic of each cMyc- or HA-tagged versions of the proteins (above 100kDa) and divided by the density of the corresponding major loading protein. Relative target protein accumulation at t0 for the CHX treatment (A,B,C and D) or no MG132 (E and F) was set to 1 and then compared across all lanes, to assess changes across samples and ARFs stability.

## DISCUSSION

AR formation is a post-embryonic process that is intrinsic to the normal development of monocots. In both monocots and dicots, it can be induced in response to diverse environmental and physiological stimuli or through horticultural practices used for vegetative propagation of many dicotyledonous species (reviewed in (Bellini et al., 2014; Steffens and Rasmussen, 2016)). Vegetative propagation is widely used in horticulture and forestry for amplification of elite genotypes obtained in breeding programs or selected from natural populations. Although this requires effective rooting of stem cuttings, this is often not achieved, and many studies conducted at physiological, biochemical and molecular levels to better understand the entire process have shown that AR formation is a heritable quantitative genetic trait controlled by multiple endogenous and environmental factors. In particular, it has been shown to be controlled by complex hormone cross-talks, in which auxin plays a central role (Lakehal and Bellini, 2019; Pacurar et al., 2014b). The specificity of auxin response is thought to depend on a specific combinatorial suite of ARF–Aux/IAA protein–protein interactions from among the huge number of potential interactions that modulate the auxin response of gene promoters via different affinities and activities (reviewed in (Vernoux et al., 2011; Weijers et al., 2005)). In previous work, we identified a regulatory module composed of three *ARF* genes, two activators (*ARF6* and *ARF8*) and one repressor (*ARF17*), which we showed could control AR formation in Arabidopsis hypocotyls (Gutierrez et al., 2009) (Supplemental Figure 1). Recent developments have highlighted the complexity of many aspects of ARF function. In particular, crystallization of the DNA binding domains of ARF1 and ARF5 (Boer et al., 2014) and the C‐terminal protein binding domain 1 (PB1) from ARF5 (Nanao et al., 2014) and ARF7 (Korasick et al., 2014) has provided insights into the physical aspects of ARF interactions and demonstrated new perspectives for dimerization and oligomerization that impact ARF functional cooperativity (Parcy et al., 2016). Here, we provide evidence that ARF6 can form a homodimer while we could detect neither heterodimerization between ARF6 and ARF8 nor ARF8 homodimerization. How this influences their respective role in the control of AR initiation is not yet known and requires further investigation. Nevertheless, based on a recent structural analysis of other ARFs (Nanao et al., 2014; Parcy et al., 2016), we propose that the ARF6 homodimer would probably target different sites from that of a monomeric ARF8 protein in the *GH3s* promotors, and/or that their respective efficiency of transcriptional regulation would be different, suggesting that one of the two transcription factors might have a prevalent role compared to the other. The prevailing model for auxin-mediated regulation of the Aux/IAA–ARF transcriptional complex is *via* increased Aux/IAA degradation in the presence of auxin, permitting ARF action, possibly through ARF-ARF dimerization, and subsequent auxin-responsive gene regulation (Nanao et al., 2014; Parcy et al., 2016). As a further step of regulation for auxin-responsive gene transcription, it has been suggested that proteasomal degradation of ARF proteins may be as important as that of Aux/IAA proteins to modulate the ratio between ARFs and Aux/IAAs proteins (Salmon et al., 2008). In the present work, we demonstrated that like ARF1 (Salmon et al., 2008), proteins ARF6, ARF8 and ARF17 undergo proteasome dependent degradation. We previously showed that the balance between the two positive regulators ARF6 and ARF8 and the negative regulator ARF17 was important for determining the number of ARs and that this balance was modulated at the post-transcriptional level by the action of the microRNAs miR167 and miR160 (Gutierrez et al., 2009). Here, we suggest that the proteasome dependent degradation of ARF6, ARF8 and ARF17 proteins is an additional level of regulation for modulation of the transcription factor balance during AR formation.

ARF6 and ARF8 (but not ARF17) retain PB1 in their structure, which makes them targets of Aux/IAA repressor proteins. Because most previous genetic studies of *Aux/IAA* genes focused on characterization of gain-of-function mutants and there are only a few recent characterizations of KO mutants (Arase et al., 2012; Shani et al., 2017), we attempted to identify potential Aux/IAA partners involved in the control of AR initiation in the Arabidopsis hypocotyl. Nevertheless, likely because AR formation is a quantitative trait, we identified six *iaa* KO mutants showing an increased number of ARs. We confirmed direct physical interaction with ARF6 and/or ARF8 for three of them (IAA6, IAA9 and IAA17) and showed significant upregulation of *GH3.3*, *GH3.5* and *GH3.6* expression in the corresponding single KO mutants, confirming that each of the three IAA proteins act as repressors in this pathway. Vernoux *et al.* (2011) also showed interaction between IAA17 and the PB1 domain of ARF6 and ARF8, but in contrast to our results, IAA9 was found to interact with ARF6 and not ARF8. The same study showed interaction of ARF6 and ARF8 with IAA7 and IAA8, which we did not observe when using the full-length proteins. Nevertheless, a KO mutation in *IAA5*, *IAA7* and *IAA8* genes led to a similar phenotype as observed in *iaa6*, *iaa9* and *iaa17* KO mutants. It is therefore possible that IAA5, IAA7 and IAA8 proteins contribute in a combinatorial manner to generate a higher order of oligomerization through interaction with one of the other three Aux/IAA proteins, leading to repression of ARF6 and ARF8 activity. Indeed, Vernoux *et al.* (2011) showed that in the yeast two-hybrid interactome, IAA5, IAA7 and IAA8 interact with IAA6, IAA9 and IAA17. Further, recent work has demonstrated that dimerization of the Aux/IAA repressor with the transcription factor is insufficient to repress the activity and that multimerization is likely to be the mechanism for repressing ARF transcriptional activity (Korasick et al., 2014), which supports our hypothesis. Alternatively, IAA5, IAA7 and IAA8 could contribute to repressing the activity of other ARFs, such as ARF7 and/or ARF19, which have also been shown to be involved in the control of AR formation (Sheng et al., 2017).

In addition to Aux/IAA transcriptional repressors and ARF transcription factors, TIR1/AFB F-box proteins are required for a proper auxin regulation of transcription. Several elegant studies have shown that auxin promotes degradation of Aux/IAA proteins through the SCF^TIR1/AFB^ in an auxin-dependent manner (Dharmasiri et al., 2005a; Gray et al., 2001; Kepinski and Leyser, 2005; Ramos et al., 2001; Tan et al., 2007)(40-44). Hence, our model would not be complete without the F-box proteins necessary to release ARF6 and ARF8 transcriptional activity. Among the six TIR1/AFB proteins examined, we demonstrated that TIR1 and AFB2 are the main players involved in this process. Both these proteins act by modulating JA homeostasis since an accumulation of JA and JA-Ile was observed in the single mutants. Nevertheless, our results suggest a different and complementary role for TIR1 and AFB2. Indeed, a mutation in the *TIR1* gene did not affect the expression of the three *GH3* genes in the same way as a mutation in the *AFB2* gene but instead mainly affected the expression of genes involved in JA biosynthesis. These results are in agreement with a previous study, which showed that TIR1 controls JA biosynthesis during flower development (Cecchetti et al., 2013). ARF6 and ARF8 have also been shown to be positive regulators of JA biosynthesis during flower development (Nagpal et al., 2005). However, it is unlikely that TIR1 controls JA biosynthesis through ARF6 and/or ARF8 during AR initiation since ARF6 and ARF8 have been shown to be positive regulators of AR initiation upstream of JA signaling (Gutierrez et al., 2009; Gutierrez et al., 2012). We are conscious that both gene expression analysis and hormone quantification were performed on whole hypocotyls, at particular time points and therefore may not fully reflect the dynamic of events in the single cells from which the AR initiate. Both gene expression analysis and hormone quantification were performed on whole hypocotyls, at particular time points and therefore may not reflect the dynamic of events in the single cells from which the AR initiate. Nevertheless, because our previous work had shown a clear correlation between *GH3* gene expression or protein content in the whole hypocotyl and the number of ARs (Pacurar et al., 2014a; Sorin et al., 2006) on a one hand, and that mutants deficient in JA biosynthesis had an increased number of ARs (Gutierrez et al., 2012) on another hand, we would like to propose here a dual role for TIR1 in the control of AR initiation, i.e., control of JA conjugation through a ARF6/ARF8 signaling module and control of JA biosynthesis through a pathway yet to be identified that would lead to similar amount of endogenous JA and JA isoleucine depending on the developmental stage.

In conclusion, we propose that AR initiation in the Arabidopsis hypocotyl depends on a regulatory module comprising two F-box proteins (TIR1 and AFB2), at least three Aux/IAA proteins (IAA6, IAA9 and IAA17) and three ARF transcriptional regulators (ARF6, ARF8 and ARF17), which control AR initiation by modulating JA homeostasis (Figure 7).

**Figure 7:**
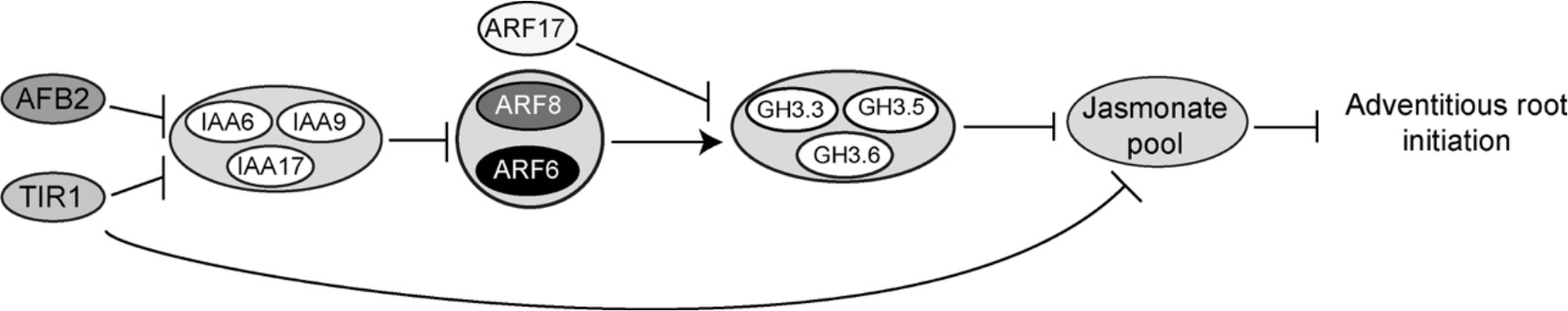
Molecular framework for TIR1/AFB-Aux/IAA-dependent auxin sensing controlling adventitious rooting in Arabidopsis. The F-box proteins TIR1 and AFB2 control JA homeostasis by promoting the degradation of IAA6, IAA9 and IAA17 protein that repress the transcriptional activity of ARF6 and ARF8. TIR1 protein has a dual role and also control JA biosynthesis through a pathway yet to be identified.

## MATERIALS AND METHODS

### Plant material and growth conditions

The single mutants *tir1-1, afb1-3, afb2-3, afb3-4, afb4-8* and *afb5-5*, multiple mutants *tir1-1afb2-3, afb2-3afb3-4, afb4-8afb5-5* and, translational fusion lines *tir1-1*pTIR1:cTIR1-GUS and *afb2-3*pAFB2:cAFB2-GUS were described in (Parry et al., 2009). Seeds of the mutants and transgenic lines were provided by Prof. Mark Estelle (UCSD, San Diego, CA, USA). The *iaa* T-DNA insertion mutants used in this study are listed in Supplemental Table 1. All the mutants were provided by the Nottingham Arabidopsis Stock Centre, except *iaa3/shy2-24*, which was provided by Prof. Jason Reed (UNC, Chapel Hill, NC, USA). The mutant lines *iaa4-1*, *iaa5-1*, *iaa6-1*, *iaa8-1*, *iaa9-1*, *iaa11-1*, *iaa12-1*, *iaa14-1*, *iaa17-6* and *iaa33-1* were previously described in (Overvoorde et al., 2005). The *Arabidopsis thaliana* ecotype Columbia-0 (Col-0) was used as the wild type and background for all the mutants and transgenic lines, except *iaa3/shy2-24*, which had a Landsberg *erecta* (L*er*) background. Growth conditions and adventitious rooting experiments were performed as previously described (Gutierrez et al., 2009; Sorin et al., 2005).

### Hormone profiling experiment

Hypocotyls from the wild type Col-0, single mutants *tir1-1* and *afb2-3* and double mutant *tir1-1afb2-3* were collected from seedlings grown as described in (Gutierrez et al., 2012). Samples were prepared from six biological replicates; for each, at least 2 technical replicates were used. Endogenous levels of free IAA, SA and JA as well as the conjugated form of JA, JA-Ile, were determined in 20 mg of hypocotyls according to the method described in (Flokova et al., 2014). The phytohormones were extracted using an aqueous solution of methanol (10% MeOH/H_2_O, v/v). To validate the LC-MS method, a cocktail of stable isotope-labeled standards was added with the following composition: 5 pmol of [^13^C_6_]IAA, 10 pmol of [^2^H_6_]JA, [^2^H_2_]JA-Ile and 20 pmol of [^2^H_4_]SA (all from Olchemim Ltd, Czech Republic) per sample. The extracts were purified using Oasis HLB columns (30 mg/1 ml, Waters) and targeted analytes were eluted using 80% MeOH. Eluent containing neutral and acidic compounds was gently evaporated to dryness under a stream of nitrogen. Separation was performed on an Acquity UPLC® System (Waters, Milford, MA, USA) equipped with an Acquity UPLC BEH C18 column (100 × 2.1 mm, 1.7 μm; Waters), and the effluent was introduced into the electrospray ion source of a triple quadrupole mass spectrometer Xevo™ TQ-S MS (Waters).

### RNA isolation and cDNA Synthesis

RNAs from the hypocotyls of Col-0 and the mutants were prepared as described by (Gutierrez et al., 2009; Gutierrez et al., 2012). The resulting RNA preparations were treated with DNaseI using a DNAfree Kit (Ambion) and cDNA was synthesized by reverse transcribing 2 μg of total RNA using SuperScript III reverse transcriptase (ThermoFisher Scientific; https://www.thermofisher.com) with 500 ng of oligo(dT)18 primer according to the manufacturer’s instructions. The reaction was stopped by incubation at 70°C for 10 min, and then the reaction mixture was treated with RNaseH (ThermoFisher Scientific; https://www.thermofisher.com) according to the manufacturer’s instructions. All cDNA samples were tested by PCR using specific primers flanking an intron sequence to confirm the absence of genomic DNA contamination.

### Quantitative RT-PCR experiments

Transcript levels were assessed in three independent biological replicates by real-time qRT-PCR), in assays with triplicate reaction mixtures (final volume 20 μl) containing 5 μl of cDNA, 0.5 μM of both forward and reverse primers and 1 × FastStart SYBR Green Master mix (Roche). Steady state levels of transcripts were quantified using primers listed in Supplemental Table 2. *APT1* and *TIP41* had previously been validated as the most stably expressed genes among 11 tested in our experimental procedures and were used to normalize the qRT-PCR data (Gutierrez et al., 2009). The normalized expression patterns obtained using the reference genes were similar. Therefore, only data normalized with *TIP41* are shown. The CT (crossing threshold value) and PCR efficiency (*E*) values were used to calculate expression using the formula 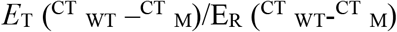, where T is the target gene, R is the reference gene, M refers to cDNA from the mutant line and WT refers to cDNA from the wild type. Data for the mutants were presented relative to those of the wild type, the calibrator.

### Heatmap of *AUXIAA* gene expression

*AUXIAA* gene expression values were obtained as described previously in different organs (cotyledons, hypocotyls and roots). The *AUXIAA* expression values for hypocotyls and roots were calculated relative to those of the cotyledon samples as calibrator and set as 1. These values were subsequently used to build a cluster heatmap using Genesis software (http://www.mybiosoftware.com/genesis-1-7-6-cluster-analysis-microarray-data.html)(Sturn et al., 2002). Genes with similar expression levels between organs were clustered based on Pearson’s correlation. Correlation values near 1 indicated a strong positive correlation between two genes.

### Tagged protein constructs

Epitope-tagged versions of ARF6, ARF8, ARF17, IAA5, IAA6, IAA7, IAA8, IAA9 and IAA17 proteins were produced in pRT104-3xHA and pRT104-3xMyc plasmids (Fulop et al., 2005). All plasmids displayed a 35S promoter sequence upstream of the multi-cloning site. The open reading frames of *ARF6*, *ARF8*, *ARF17*, *IAA5*, *IAA6*, *IAA7*, *IAA8*, *IAA9* and *IAA17* were amplified from cDNA from 7-day-old *Arabidopsis* seedlings using Finnzyme’s Phusion high-fidelity DNA polymerase protocol with gene-specific primers listed in *SI Appendix* Table S3. For the bimolecular functional complementation assay (BiFC), the open reading frames of *ARF6*, *ARF8*, *IAA6*, *IAA9* and *IAA17* were amplified with gene-specific primers carrying BgIII or KpnI restriction sites to facilitate subsequent cloning (*SI Appendix* Table S4). The products obtained after PCR were digested with BgIII and KpnI prior to ligation into pSAT-nEYFP and pSAT-cEYFP plasmids (Citovsky et al., 2006) that had previously been cut open with the same enzymes. All constructs were verified by sequencing.

### Protoplast production and transformation

Protoplasts from *Arabidopsis* cell culture or 14-day-old Arabidopsis seedlings were prepared and transfected as previously described (Meskiene et al., 2003; Zhai et al., 2009). For CoIP, 10^5^ protoplasts from the Arabidopsis cell culture were transfected with 5 to 7.5 μg of each construct. For BiFC assays, Arabidopsis mesophyll protoplasts were co-transfected with 10 µg of each construct. The protoplasts were imaged by confocal laser scanning microscopy after 24 hours of incubation in the dark at room temperature.

### Co-immunoprecipitation

For testing protein interactions, co-transfected protoplasts were extracted in lysis buffer containing 25 mM Tris-HCl, pH 7.8, 10 mM MgCl_2_, 75 mM NaCl, 5 mM EGTA, 60 mM β-glycerophosphate, 1 mM dithiothreitol, 10% glycerol, 0.2% Igepal CA-630 and Protein Inhibitor Cocktail (Sigma-Aldrich; http://www.sigmaaldrich.com/). The cell suspension was frozen in liquid nitrogen and then thawed on ice and centrifuged for 5 min at 150 g. The resulting supernatant was mixed with 1.5 μl of anti-Myc antibody (9E10, Covance; http://www.covance.com/) or 2 μl of anti-HA antibody (16B12, Covance; http://www.covance.com/)] for 2 h at 4°C on a rotating wheel. Immunocomplexes were captured on 10 μl of Protein G-Sepharose beads, washed three times in 25 mM sodium phosphate, 5% glycerol and 0.2% Igepal CA-630 buffer and then eluted by boiling with 40 μl of SDS sample buffer. The presence of immunocomplexes was assessed by probing protein gel blots with either anti-HA (3F10, Sigma/Roche; http://www.sigmaaldrich.com/) or anti-Myc antibody (9E10, Covance; http://www.covance.com/) at 1:2000 dilution.

### Cycloheximide or proteasome inhibitor treatment of transfected protoplasts

Sixteen hours after protoplast transfection, cycloheximide (CHX) (SigmaAldrich; http://www.sigmaaldrich.com/) was added to a final concentration of 200 μg/ml in the protoplast growth medium and the protoplasts were incubated for 0, 0.5, 1, 1.5 and 2 h. Afterwards, the protoplasts were harvested and the proteins extracted and analyzed by SDS-PAGE and western blotting.

The proteasome inhibitor MG132 (SigmaAldrich; http://www.sigmaaldrich.com/) was applied at a concentration of 50 µM 16 h after protoplasts transfection. After 2 h incubation, the protoplasts were harvested and the proteins were extracted and analyzed by SDS-PAGE and western blotting. The plasmid expressing *HA_3_-ARF1* was described in (Salmon et al., 2008) and kindly provided by Prof. Judy Callis (UC, Davis, CA, USA).

### Proteasome inhibition in planta

Seeds from Arabidopsis lines expressing HA_3_:ARF1, cMyc_3_:ARF6, cMyc_3_:ARF8 and cMyc_3_:ARF17 were sterilized and sown *in vitro* as previously described (Sorin et al., 2005). Plates were incubated at 4°C for 48 h for stratification and transferred to the light for 16 h at a temperature of 20°C to induce germination. The plates were then wrapped in aluminum foil and kept until the hypocotyl of the seedlings reached on average 6 mm. The plates were then transferred back to the light for 6 days. On day 6, the seedlings were transferred to liquid growth medium (GM). On day 7, the GM was removed and fresh GM without (DMSO control) or with MG132 (SigmaAldrich, http://www.sigmaaldrich.com/) at a final concentration of 100 μM was added, and the seedlings incubated for a further 2 h. After incubation, the GM liquid culture was removed, and proteins were extracted and analyzed by SDS-PAGE and western blotting. The Arabidopsis line expressing *HA_3_-ARF1* was described in (Salmon et al., 2008) and kindly provided by Prof. Judy Callis (UC, Davis, CA, USA).

### Analysis of promoter activity

A 1-kb-long fragment upstream from the start codon of *IAA6*, *IAA9* and *IAA17* was amplified by applying PCR to Col-0 genomic DNA. The primer sequences used are listed in *SI Appendix* Table S5. The amplified fragments were cloned using a pENTR/D-TOPO cloning kit (ThermoFisher Scientific; https://www.thermofisher.com) and transferred into the pKGWFS7 binary vector (Karimi et al., 2002) using a Gateway LR Clonase enzyme mix (ThermoFisher Scientific; https://www.thermofisher.com) according to the manufacturer’s instructions. Transgenic Arabidopsis plants expressing the *promIAA6:GUS*, *promIAA9:GUS* and *promIAA17:GUS* fusion were generated by *Agrobacterium tumefaciens* mediated floral dipping and the expression pattern was checked in the T2 progeny of several independent transgenic lines. Histochemical assays of GUS expression were performed as previously described (Sorin et al., 2005).

### Confocal laser scanning microscopy

For the BIFC assay, images of fluorescent protoplasts were obtained with a Leica TCS-SP2-AOBS spectral confocal laser scanning microscope equipped with a Leica HC PL APO × 20 water immersion objective. YFP and chloroplasts were excited with the 488 nm line of an argon laser (laser power 35%). Fluorescence emission was detected over the range 495 to 595 nm for the YFP construct and 670 to 730 nm for chloroplast autofluorescence. Images were recorded and processed using LCS software version 2.5 (Leica Microsytems). Images were cropped using Adobe Photoshop CS2 and assembled using Adobe Illustrator CS2 software (Abode, http://www.abode.com).

## ACKNOWLEDGMENTS

The authors would like to thank Prof. Mark Estelle (UCSD, San Diego, CA, USA) and Prof Jason Reed (UNC, Chapel Hill, NC, USA) for providing seeds of single and multiple mutants. The authors also thank Prof. Judy Callis (UC, Davis, CA, USA) for providing Arabidopsis line and the plasmid expressing *HA_3_-ARF1.* We also thank Hana Martinková for help with phytohormone analyses. This work was supported by the Swedish Research Council (VR), the Swedish Research Council for Research and Innovation for Sustainable Growth (VINNOVA), the K. & A. Wallenberg Foundation, the Carl Trygger Foundation, the Carl Kempe Foundation, the University of Picardie Jules Verne, the Regional Council of Picardie, the European Regional Development Fund, and the Ministry of Education, Youth and Sports of the Czech Republic (European Regional Development Fund-Project “Plants as a tool for sustainable global development” no. CZ.02.1.01/0.0/0.0/16_019/0000827).

## AUTHORS CONTRIBUTION

A.L. and S.C. contributed equally to this work.

A.L., S.C., E.C., R.L.H., Z.R., O.N., F.J., D.I.P., I.P., A.R., L.G., L.B. performed or contributed to the experiments. C.B., A.L., S.C. and E.C. designed the research and analyzed the data. C.B. conceptualized and supervised the overall project. C.B. wrote the article with input from A.L.. All authors read and commented on the manuscript.

## REFERENCES

Arase, F., Nishitani, H., Egusa, M., Nishimoto, N., Sakurai, S., Sakamoto, N., and Kaminaka, H. (2012). IAA8 involved in lateral root formation interacts with the TIR1 auxin receptor and ARF transcription factors in Arabidopsis. PLoS One 7:e43414.

Bellini, C., Pacurar, D.I., and Perrone, I. (2014). Adventitious roots and lateral roots: similarities and differences. Annu. Rev. Plant Biol. 65:639–666.

Boer, D.R., Freire-Rios, A., van den Berg, W.A., Saaki, T., Manfield, I.W., Kepinski, S., Lopez-Vidrieo, I., Franco-Zorrilla, J.M., de Vries, S.C., Solano, R., et al. (2014). Structural basis for DNA binding specificity by the auxin-dependent ARF transcription factors. Cell 156:577–589.

Calderon Villalobos, L.I., Lee, S., De Oliveira, C., Ivetac, A., Brandt, W., Armitage, L., Sheard, L.B., Tan, X., Parry, G., Mao, H., et al. (2012). A combinatorial TIR1/AFB-Aux/IAA co-receptor system for differential sensing of auxin. Nat. Chem. Biol. 8:477–485.

Cecchetti, V., Altamura, M.M., Brunetti, P., Petrocelli, V., Falasca, G., Ljung, K., Costantino, P., and Cardarelli, M. (2013). Auxin controls Arabidopsis anther dehiscence by regulating endothecium lignification and jasmonic acid biosynthesis. Plant J. 74:411–422.

Chapman, E.J., and Estelle, M. (2009). Mechanism of auxin-regulated gene expression in plants. Annu. Rev. Genet. 43:265–285.

Citovsky, V., Lee, L.Y., Vyas, S., Glick, E., Chen, M.H., Vainstein, A., Gafni, Y., Gelvin, S.B., and Tzfira, T. (2006). Subcellular localization of interacting proteins by bimolecular fluorescence complementation in planta. J. Mol. Biol. 362:1120–1131.

De Rybel, B., Vassileva, V., Parizot, B., Demeulenaere, M., Grunewald, W., Audenaert, D., Van Campenhout, J., Overvoorde, P., Jansen, L., Vanneste, S., et al. (2010). A novel aux/IAA28 signaling cascade activates GATA23-dependent specification of lateral root founder cell identity. Curr. Biol. 20:1697–1706.

De Smet, I., Lau, S., Voss, U., Vanneste, S., Benjamins, R., Rademacher, E.H., Schlereth, A., De Rybel, B., Vassileva, V., Grunewald, W., et al. (2010). Bimodular auxin response controls organogenesis in Arabidopsis. Proc. Natl. Acad. Sci. U S A 107:2705–2710.

Dello Ioio, R., Nakamura, K., Moubayidin, L., Perilli, S., Taniguchi, M., Morita, M.T., Aoyama, T., Costantino, P., and Sabatini, S. (2008). A genetic framework for the control of cell division and differentiation in the root meristem. Science 322:1380–1384.

Dharmasiri, N., Dharmasiri, S., and Estelle, M. (2005a). The F-box protein TIR1 is an auxin receptor. Nature 435:441–445.

Dharmasiri, N., Dharmasiri, S., Weijers, D., Lechner, E., Yamada, M., Hobbie, L., Ehrismann, J.S., Jurgens, G., and Estelle, M. (2005b). Plant development is regulated by a family of auxin receptor F box proteins. Dev. Cell. 9:109–119.

Flokova, K., Tarkowska, D., Miersch, O., Strnad, M., Wasternack, C., and Novak, O. (2014). UHPLC-MS/MS based target profiling of stress-induced phytohormones. Phytochemistry 105:147–157.

Fukaki, H., Tameda, S., Masuda, H., and Tasaka, M. (2002). Lateral root formation is blocked by a gain-of-function mutation in the SOLITARY-ROOT/IAA14 gene of Arabidopsis. Plant J. 29:153–168.

Fulop, K., Pettko-Szandtner, A., Magyar, Z., Miskolczi, P., Kondorosi, E., Dudits, D., and Bako, L. (2005). The Medicago CDKC;1-CYCLINT;1 kinase complex phosphorylates the carboxy-terminal domain of RNA polymerase II and promotes transcription. Plant J. 42:810–820.

Geiss, G., Gutierrez, L., and Bellini, C. (2009). Adventitious root formation: new insights and perspective. In: Root Development - Annual Plant Reviews --Beeckman, T., ed. London: A John Wiley & Sons, Ltd. 127–156.

Gray, W.M., Kepinski, S., Rouse, D., Leyser, O., and Estelle, M. (2001). Auxin regulates SCF(TIR1)-dependent degradation of AUX/IAA proteins. Nature 414:271–276.

Guilfoyle, T.J., and Hagen, G. (2007). Auxin response factors. Curr. Opin. Plant Biol. 10:453–460.

Guilfoyle, T.J., and Hagen, G. (2012). Getting a grasp on domain III/IV responsible for Auxin Response Factor-IAA protein interactions. Plant Sci. 190:82–88.

Gutierrez, L., Bussell, J.D., Pacurar, D.I., Schwambach, J., Pacurar, M., and Bellini, C. (2009). Phenotypic plasticity of adventitious rooting in Arabidopsis is controlled by complex regulation of AUXIN RESPONSE FACTOR transcripts and microRNA abundance. Plant Cell 21:3119–3132.

Gutierrez, L., Mongelard, G., Flokova, K., Pacurar, D.I., Novak, O., Staswick, P., Kowalczyk, M., Pacurar, M., Demailly, H., Geiss, G., et al. (2012). Auxin controls Arabidopsis adventitious root initiation by regulating jasmonic acid homeostasis. Plant Cell 24:2515–2527.

Hamann, T., Benkova, E., Baurle, I., Kientz, M., and Jurgens, G. (2002). The Arabidopsis BODENLOS gene encodes an auxin response protein inhibiting MONOPTEROS-mediated embryo patterning. Genes Dev. 16:1610–1615.

Karimi, M., Inze, D., and Depicker, A. (2002). GATEWAY vectors for Agrobacterium-mediated plant transformation. Trends Plant Sci. 7:193–195.

Kepinski, S., and Leyser, O. (2005). The Arabidopsis F-box protein TIR1 is an auxin receptor. Nature 435:446–451.

Korasick, D.A., Westfall, C.S., Lee, S.G., Nanao, M.H., Dumas, R., Hagen, G., Guilfoyle, T.J., Jez, J.M., and Strader, L.C. (2014). Molecular basis for AUXIN RESPONSE FACTOR protein interaction and the control of auxin response repression. Proc. Natl. Acad. Sci. U S A 111:5427–5432.

Lakehal, A., and Bellini, C. (2019). Control of adventitious root formation: insights into synergistic and antagonistic hormonal interactions. Physiol Plant. 165:90–100.

Lavenus, J., Goh, T., Roberts, I., Guyomarc’h, S., Lucas, M., De Smet, I., Fukaki, H., Beeckman, T., Bennett, M., and Laplaze, L. (2013). Lateral root development in Arabidopsis: fifty shades of auxin. Trends Plant Sci. 18:450–458.

Magyar, Z., De Veylder, L., Atanassova, A., Bako, L., Inze, D., and Bogre, L. (2005). The role of the Arabidopsis E2FB transcription factor in regulating auxin-dependent cell division. Plant Cell 17:2527–2541.

Meskiene, I., Baudouin, E., Schweighofer, A., Liwosz, A., Jonak, C., Rodriguez, P.L., Jelinek, H., and Hirt, H. (2003). Stress-induced protein phosphatase 2C is a negative regulator of a mitogen-activated protein kinase. J. Biol. Chem. 278:18945–18952.

Nagpal, P., Ellis, C.M., Weber, H., Ploense, S.E., Barkawi, L.S., Guilfoyle, T.J., Hagen, G., Alonso, J.M., Cohen, J.D., Farmer, E.E., et al. (2005). Auxin response factors ARF6 and ARF8 promote jasmonic acid production and flower maturation. Development 132:4107–4118.

Nanao, M.H., Vinos-Poyo, T., Brunoud, G., Thevenon, E., Mazzoleni, M., Mast, D., Laine, S., Wang, S., Hagen, G., Li, H., et al. (2014). Structural basis for oligomerization of auxin transcriptional regulators. Nat. Commun. 5:3617.

Orosa-Puente, B., Leftley, N., von Wangenheim, D., Banda, J., Srivastava, A.K., Hill, K., Truskina, J., Bhosale, R., Morris, E., Srivastava, M., et al. (2018). Root branching toward water involves posttranslational modification of transcription factor ARF7. Science 362:1407–1410.

Overvoorde, P.J., Okushima, Y., Alonso, J.M., Chan, A., Chang, C., Ecker, J.R., Hughes, B., Liu, A., Onodera, C., Quach, H., et al. (2005). Functional genomic analysis of the AUXIN/INDOLE-3-ACETIC ACID gene family members in Arabidopsis thaliana. Plant Cell 17:3282–3300.

Pacurar, D.I., Pacurar, M.L., Bussell, J.D., Schwambach, J., Pop, T.I., Kowalczyk, M., Gutierrez, L., Cavel, E., Chaabouni, S., Ljung, K., et al. (2014a). Identification of new adventitious rooting mutants amongst suppressors of the Arabidopsis thaliana superroot2 mutation. J Exp Bot 65:1605–1618.

Pacurar, D.I., Perrone, I., and Bellini, C. (2014b). Auxin is a central player in the hormone cross-talks that control adventitious rooting. Physiol. Plant. 151:83–96.

Parcy, F., Vernoux, T., and Dumas, R. (2016). A Glimpse beyond Structures in Auxin-Dependent Transcription. Trends Plant Sci. 21:574–583.

Parry, G., Calderon-Villalobos, L.I., Prigge, M., Peret, B., Dharmasiri, S., Itoh, H., Lechner, E., Gray, W.M., Bennett, M., and Estelle, M. (2009). Complex regulation of the TIR1/AFB family of auxin receptors. Proc. Natl. Acad. Sci. U S A 106:22540–22545.

Ramos, J.A., Zenser, N., Leyser, O., and Callis, J. (2001). Rapid degradation of auxin/indoleacetic acid proteins requires conserved amino acids of domain II and is proteasome dependent. Plant Cell 13:2349–2360.

Salmon, J., Ramos, J., and Callis, J. (2008). Degradation of the auxin response factor ARF1. Plant J. 54:118–128.

Shani, E., Salehin, M., Zhang, Y., Sanchez, S.E., Doherty, C., Wang, R., Mangado, C.C., Song, L., Tal, I., Pisanty, O., et al. (2017). Plant Stress Tolerance Requires Auxin-Sensitive Aux/IAA Transcriptional Repressors. Curr. Biol. 27:437–444.

Sheng, L., Hu, X., Du, Y., Zhang, G., Huang, H., Scheres, B., and Xu, L. (2017). Non-canonical WOX11-mediated root branching contributes to plasticity in Arabidopsis root system architecture. Development 144:3126–3133.

Sorin, C., Bussell, J.D., Camus, I., Ljung, K., Kowalczyk, M., Geiss, G., McKhann, H., Garcion, C., Vaucheret, H., Sandberg, G., et al. (2005). Auxin and light control of adventitious rooting in Arabidopsis require ARGONAUTE1. Plant Cell 17:1343–1359.

Sorin, C., Negroni, L., Balliau, T., Corti, H., Jacquemot, M.P., Davanture, M., Sandberg, G., Zivy, M., and Bellini, C. (2006). Proteomic analysis of different mutant genotypes of Arabidopsis led to the identification of 11 proteins correlating with adventitious root development. Plant Physiol 140:349–364.

Steffens, B., and Rasmussen, A. (2016). The Physiology of Adventitious Roots. Plant Physiol 170:603–617.

Sturn, A., Quackenbush, J., and Trajanoski, Z. (2002). Genesis: cluster analysis of microarray data. Bioinformatics 18:207–208.

Sun, J., Qi, L., Li, Y., Zhai, Q., and Li, C. (2013). PIF4 and PIF5 transcription factors link blue light and auxin to regulate the phototropic response in Arabidopsis. Plant Cell 25:2102–2114.

Szemenyei, H., Hannon, M., and Long, J.A. (2008). TOPLESS mediates auxin-dependent transcriptional repression during Arabidopsis embryogenesis. Science 319:1384–1386.

Tan, X., Calderon-Villalobos, L.I., Sharon, M., Zheng, C., Robinson, C.V., Estelle, M., and Zheng, N. (2007). Mechanism of auxin perception by the TIR1 ubiquitin ligase. Nature 446:640–645.

Tatematsu, K., Kumagai, S., Muto, H., Sato, A., Watahiki, M.K., Harper, R.M., Liscum, E., and Yamamoto, K.T. (2004). MASSUGU2 encodes Aux/IAA19, an auxin-regulated protein that functions together with the transcriptional activator NPH4/ARF7 to regulate differential growth responses of hypocotyl and formation of lateral roots in Arabidopsis thaliana. Plant Cell 16:379–393.

Vernoux, T., Brunoud, G., Farcot, E., Morin, V., Van den Daele, H., Legrand, J., Oliva, M., Das, P., Larrieu, A., Wells, D., et al. (2011). The auxin signalling network translates dynamic input into robust patterning at the shoot apex. Mol. Syst. Biol. 7:508.

Wang, H., Jones, B., Li, Z., Frasse, P., Delalande, C., Regad, F., Chaabouni, S., Latche, A., Pech, J.C., and Bouzayen, M. (2005). The tomato Aux/IAA transcription factor IAA9 is involved in fruit development and leaf morphogenesis. Plant Cell 17:2676–2692.

Wang, R., and Estelle, M. (2014). Diversity and specificity: auxin perception and signaling through the TIR1/AFB pathway. Curr. Opin. Plant Biol. 21:51–58.

Weijers, D., Sauer, M., Meurette, O., Friml, J., Ljung, K., Sandberg, G., Hooykaas, P., and Offringa, R. (2005). Maintenance of embryonic auxin distribution for apical-basal patterning by PIN-FORMED-dependent auxin transport in Arabidopsis. Plant Cell 17:2517–2526.

Weijers, D., and Wagner, D. (2016). Transcriptional Responses to the Auxin Hormone. Annu. Rev. Plant Biol. 67:539–574.

Zhai, Z., Jung, H.I., and Vatamaniuk, O.K. (2009). Isolation of protoplasts from tissues of 14-day-old seedlings of Arabidopsis thaliana. J. Vis. Exp. 30:1149

