## Supplemental Information for "Molecular framework for TIR1/AFB-Aux/IAA-dependent auxin sensing controlling adventitious rooting in Arabidopsis"

**This PDF includes:**

Supplemental Figures 1 to 3

Supplemental Tables 1 to 5

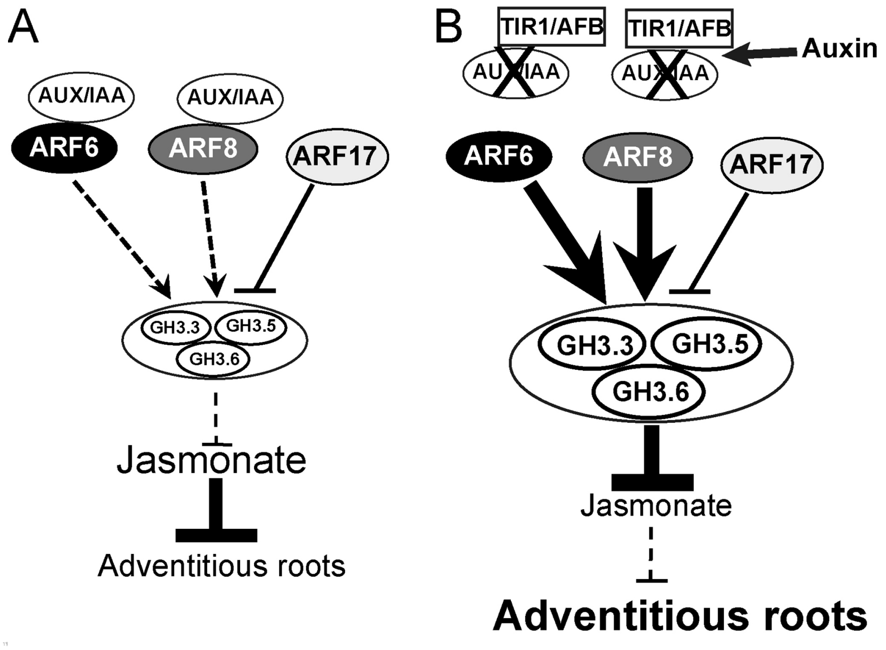


**Supplemental Figure 1: Model for regulation of adventitious root initiation by auxin**
Adventitious root initiation is controlled by a subtle balance of activator and repressor *ARF* transcripts acting upstream of JA signaling (Gutierrez et al., 2012). *ARF6* and *ARF8* are positive regulators, whereas *ARF17* is a negative regulator. Under steady-state conditions, the transcriptional activity of ARF6 and ARF8 proteins is negatively regulated by interaction with Aux/IAA proteins. This is not the case for ARF17, which lacks the PBI domain (A). Instead, the balance between positive and negative regulators leads to a steady-state AR phenotype. When auxin is added (B), the Aux/IAA proteins form an auxin coreceptor complex with auxin F-box proteins (TIR/AFB) and are sent for degradation through the 26S proteasome. In this case, the transcriptional activity of ARF6 and ARF8 is released and they induce expression of three GH3 genes that contribute to downregulating JA signaling, resulting in increased AR initiation (B).

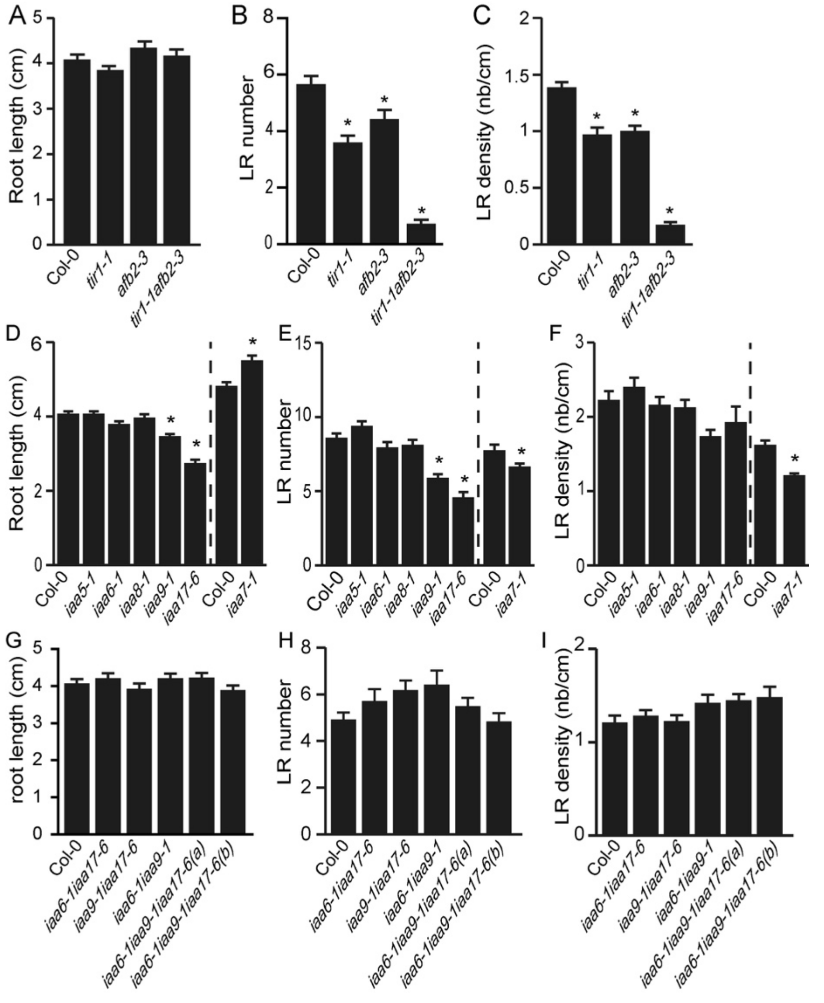


**Supplemental Figure 2: Root length, lateral root number and lateral root density of *tir/afb* and *iaa* mutants**

The primary root length (A, D, G) and number of lateral roots (LRs) (B, E, H) were measured and counted on seedlings that were first etiolated in the dark until their hypocotyls had reached 6 mm long and then transferred to the light for 7 days. (C, F, I) The mean LR density was expressed as the number of LRs divided by the length of the main root of at least 30 seedlings for each line. The LR density was calculated as the number of LRs per cm of primary root length.

Error bars= +/- SEM. One-way ANOVA combined with Dunnett's multiple comparison test indicated that some values (indicated by *) were significantly different from wild-type values.

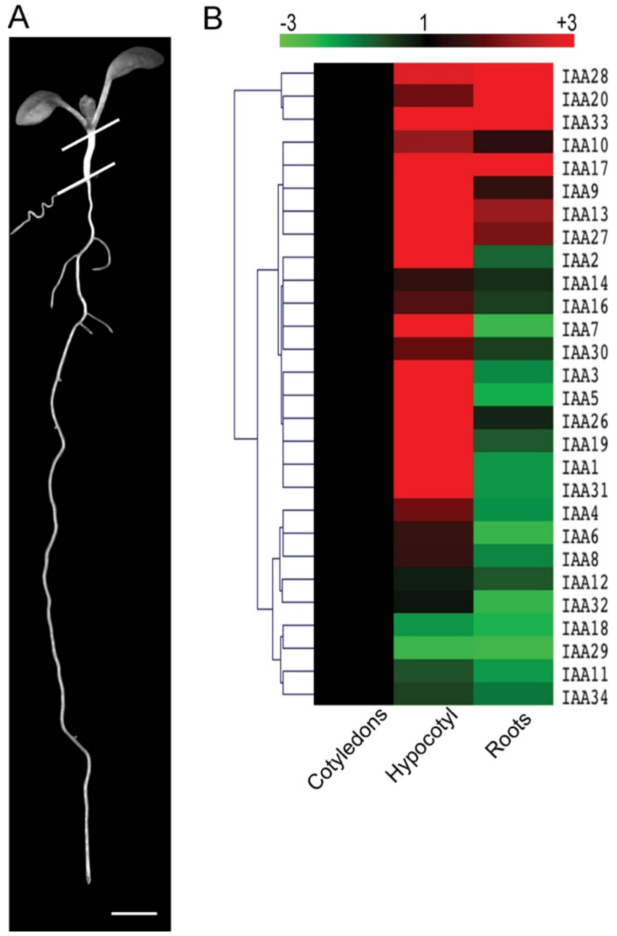


**Supplemental Figure 3: Relative expression of IAA genes in cotyledons, the hypocotyl and roots from 7-day-old Arabidopsis seedlings**

(A) Photograph of a Col-0 7-day-old seedling grown under long-day conditions illustrating the material used for quantitative RT-PCR performed in (B)

(B) Cluster heatmap of *IAA* gene expression. *IAA* gene expression values were obtained as described in the Materials and Methods section in different organs (cotyledons, hypocotyls and roots). The *IAA* gene expression values for hypocotyls and roots were calculated relative to the cotyledon samples as calibrator and set as 1.

These values were then used to build a cluster heatmap using Genesis software (http://genome.tugraz.at/genesisclient/genesisclient_description.shtml). Genes with similar expression levels between organs were clustered based on Pearson's correlation. Correlation values near 1 indicate a strong positive correlation between two genes.

**Supplemental Table 1:**

Summary of *aux/iaa* T-DNA insertion mutants used in this study

| Gene | Gene number | Allele name | NASC ID | Collection | Reference |
|---|---|---|---|---|---|
| *IAA3/SHY2* | At1g04240 | *shy2-24* | | | (Tian et al., 2002) |
| *IAA4* | At5g43700 | *iaa4-1* | N25208 | SALK | (Overvoorde et al., 2005) |
| *IAA5* | At1g15580 | *iaa5-1* | N9578 | SALK | (Alonso et al., 2003) |
| *IAA6* | At1g52830 | *iaa6-1* | N25209 | SALK | (Overvoorde et al., 2005) |
| | | *iaa6-2* | N877906 | SAIL_905_C12 | This article |
| *IAA7/AXR2* | At3g23050 | *iaa7-1* | N678809 | SALK_089809 | This article |
| *IAA8* | At2g22670 | *iaa8-1* | N25210 | SALK | (Overvoorde et al., 2005) |
| *IAA9* | At5g65670 | *iaa9-1* | N25211 | SALK | (Overvoorde et al., 2005) |
| | | *iaa9-2* | N658711 | SALK_057396C | This article |
| *IAA11* | At4g28640 | *iaa11-1* | N25212 | SALK | (Overvoorde et al., 2005) |
| *IAA12/BDL* | At1g04550 | *iaa12-1* | N25213 | SALK | (Overvoorde et al., 2005) |
| *IAA14/SLR* | At4g14550 | *iaa14-1* | N25214 | SALK | (Overvoorde et al., 2005) |
| *IAA17/AXR3* | At1g04250 | *iaa17-6* | N25216 | SALK | (Overvoorde et al., 2005) |
| | | *iaa17-1* | N658276 | SALK_065697C | (Shi et al., 2015) |
| | | *iaa17-2* | N660304 | SALK_011820 | (Shi et al., 2015) |
| *IAA28* | At5g25890 | *iaa28-1* | N669043 | Salk_129988C | (Alonso et al., 2003) |
| *IAA29* | At4g32280 | *iaa29-1* | N663323 | Salk_091933C | (Alonso et al., 2003) |
| *IAA30* | At3g62100 | *iaa30-2* | N668427 | Salk_065384C | (Alonso et al., 2003) |
| *IAA31* | At3g17600 | *iaa31-1* | N25218 | SALK | (Overvoorde et al., 2005) |
| *IAA33* | At5g57420 | *iaa33-1* | N31388 | SALK | (Overvoorde et al., 2005) |

### Supplemental Table 2: Primers used for quantifying genes by RT-PCR

| Gene name | Gene number | Forward primer | Reverse primer |
|---|---|---|---|
| *IAA1* | At4g14560 | CGGTTAGATCTCACTGGAGGCCAT | ATCTGCTCCTCCTCCTGCAAAAAC |
| *IAA2* | At3g23030 | ACCTCCTACCAAAACTCAAATCGT | GCTCGGGGTAGTTTTTGTATGTCT |
| *IAA3* | At1g04240 | TCGGGCAAGATCTATGTTCA | ACCTTTTGCCCTGTTTCTGA |
| *IAA4* | At5g43700 | TGGGATTACCAGGGACAGAA | TCTGAGCCTTTGGAGGAGAA |
| *IAA5* | At1g15580 | TTCCGCTCTGCAAATTCTGTTCG | CGATCCAAGGAACATTTCCCAAGG |
| *IAA6* | At1g52830 | TGCCAAGGTACATCTCCGACGA | TAGGAGTGGCGAAGGAGGGTAAGA |
| *IAA7* | At3g23050 | GCCATCCCACCACTTGTGCTTTAG | TCTGCTGTTCCCAAGGAGAAGACT |
| *IAA8* | At2g22670 | GCCAAGGCACAGGTTGTTGGTT | TCCATGCTCACCTTCACAAACAGA |
| *IAA9* | At5g65670 | GCCTCCCATCAACTTCGTCACTGT | CAGCCAAGGCACAAATTGTCGG |
| *IAA10* | At1g04100 | AGCCGCCTTCCGTAGCTAAAGACT | CGCAACCAGACAAGTTGCTGTAGG |
| *IAA11* | At4g28640 | CAACTAGTGGGCAAGTTGTGGGAT | TCAGTGGCTGAAGCCTTAGCTTGG |
| *IAA12* | At1g04550 | TGGGTCTAAACGCTCTGCTGAATC | ACCACTTGACTTGAACGAGGAGGA |
| *IAA13* | At2g33310 | GCTAATGGACTCGCTGCACGAAAT | TAAACCGGCTGCTTTCGCTGTCTC |
| *IAA14* | At4g14550 | ATGACTCGACAAACATCGGCCAGG | ACGAGGACAAAGATGGTGACTGGA |
| *IAA15* | At1g80390 | CCACAAACATCATCCACGGCACAT | GGAGAGGAAGGGAGAGTTTGTTGC |
| *IAA16* | At3g04730 | CGGACATGACGTTCTTGCGGAAAG | AAACCACCAGCCAAGGCACAA |
| *IAA17* | At1g04250 | GGAGCACCGTACTTGAGGAA | TTTGCCCATGGTAAAAGAGC |
| *IAA18* | At1g51950 | CAGAACCAAAGAGACAAGGAGGCA | TTTGAGCTGCAAGAAGACCTCTGA |
| *IAA19* | At3g15540 | TCGGTGTGGCCTTGAAAGATGG | TGCATGACTCTAGAAACATCCCCC |
| *IAA20* | At2g46990 | CGCATCCATTCTCTGGGCTGAAGA | TCTCCAACCATCATCCAGTCACCT |
| *IAA26* | At3g16500 | CAATGAAGGGGACAAGATGC | CCAAATGTCAAGGCAGATGA |
| *IAA27* | At4g29080 | CCATGGAAGCAACAACAATG | AACCGGGTAAACCGAGTCTT |
| *IAA28* | At5g25890 | GCTCCTCCTTGTCACCAATTCACT | ACTGGAGCTACCTCAACCCTGTTA |
| *IAA29* | At4g32280 | AAGATGGATGGTGTGGCAAT | ATCCCCTCGAAGTAGCCAGT |
| *IAA30* | At3g62100 | TGCTTCAATCCTTTGGGCTGAAGA | TCTCCAACCATCATCCAGTCACCT |
| *IAA31* | At3g17600 | TCCGACCATCATCCAATCTCCATC | TGCGGTAATCGAGATCGAAAACAT |
| *IAA32* | At2g01200 | AAGCATCAATGGACCCAAAC | GATTACCCCACCACCCTTTT |
| *IAA33* | At5g57420 | TGTGTTTCCCTTGACCGGCAAGAT | TGGAAGGACTTTGTTCGTGTAGCG |
| *IAA34* | At1g15050 | CGCAAGGTCTGTGTTCTTGA | CCCAAACATGTCCTCGAGTT |
| *ARF6* | At1g30330 | CAAAGTTTAGCAGCTACCACGA | ACGTCGTTCTCTCGGTCAAC |
| *ARF8* | At5g37020 | TTTGCTATCGAAGGGTTGTTG | CATGGGTCATCACCAAGGA |
| *ARF17* | At1g77850 | GCACCTGATCCAAGTCCTTC | GGTGAATAGCTGGGGAGGAT |
| *GH3.3* | At1g77850 | ACAATTCCGCTCCACAGTTC | ACGAGTTCCTTGCTCTCCAA |
| *GH3.5* | At4g27260 | GTCTTCGAGGACTGCTGCTT | ATGTCCCTGGCTCAACAATC |
| *GH3.6* | At5g54510 | CCTTGTTCCGTTTGATGCTT | CGTGTTACCGTTCAAGCAGA |
| *GH3.10* | At4g03400 | GGGAAATCAGAGGAGAAGCA | AACGTTCCTCGTTCCACAAC |
| *GH3.11* | At2g39940 | GTTCATCGGCTGGACAGTTT | TCAAAACGCTGTGCTGAAGT |
| *MYC2* | At1G32640 | GTGCGGGATTAGCTGGTAAA | ATGCATCCCAAACACTCCTC |
| *MYC3* | At5G46760 | TGTTGAAGCAGAGAGGCAGA | CTCCGAGAAGCGAAGCTTTA |
| *MYC4* | At4G17880 | AGGAGCAAACGAGAACTGGA | CCATCTCCCCAACCTAACAA |
| *OPR3* | At2G06050 | TGGTTGGCATGCTCAATAAG | GCCTTCCAGACTCTGTTTGC |
| *OPLC1* | At1G20510 | GGGTTATCAGGTTGCTCCAGCTGAG | CCCCACTTCTTTGTCCGGAAACGGG |
| *LOX2* | At3G45140 | GGACTCATGCCTGTACGGAGCC | TGGCGTGCACGAGCGTTGAT |
| *AOC1* | At3G25760 | CTCTCAGAACTTGGGAAATACCGA | AATGGGACGAGATCTCCGAGA |
| *AOC2* | At3G25770 | GGTGCCTACGGACAGGTCAAGC | GCGGTACCGGTGTTCCGGTG |
| *AOC3* | AT3G25780 | CGAAGGAGATAGAAACAGTCCAGC | CCGAGACAAAGCTCTGTTGGTT |
| *AOC4* | AT1G13280 | GCCGTTCTCGTAAGCGTAATGT | GGAGTTCACGCGCTTAAATCC |
| *TIP41* | At4g34270 | GCTCATCGGTACGCTCTTTT | TCCATCAGTCAGAGGCTTCC |
| *APT1* | At1g27450 | GAGACATTTTGCGTGGGATT | CGGGGATTTTAAGTGGAACA |

**Supplemental Table 3:**

Primers used for amplifying open reading frames of *ARF6*, *ARF8*, *IAA5*, *IAA6*, *IAA7*, *IAA8*, *IAA9* and *IAA17* for cloning into pRT104-3xHA and pRT104-3xMyc plasmids for subsequent CoIP assay

| | **Forward primer** | **Reverse primer** |
|---|---|---|
| *ARF6* | ATCGGAATTCATGAGATTATCTTCAGCT | CGATGGTACCCTAGTAGTTGAATGAACC |
| *ARF8* | CCCGGGATGAAGCTGTCAACATCTGG | GTCGACCTAGAGATGGGTCGGGTTTTGC |
| *IAA5* | GGATCCATGGCGAATGAGAGTAATAATC | GTCGACTCATCCTCTGTTACATGATCTC |
| *IAA6* | GGATCCATGGCAAAGGAAGGTCTAG | GTCGACTTAATCTTGCTGGAGACC |
| *IAA7* | GCGCGCGAATTCATGATCGGCCAACTTATG | GCGCGCGTCGACTCAAGATCTGTTCTTGC |
| *IAA8* | GGATCCATGTCTTATCGATTGCTAAGTG | GGTACCTCAAACCCGCTCTTTGTTCTTC |
| *IAA9* | GCGCGCGAATTCATGTCCCCGGAAGAGG | GCGCGCGTCGACTTAAGCTCTCATCTTCG |
| *IAA17* | GGATCCATGATGGGCAGTGTCGAGC | GTCGACTCAAGCTCTGCTCTTGCAC |

**Supplemental Table 4:**

Primers used for amplifying open reading frames of *ARF6*, *ARF8*, *IAA6*, *IAA9* and *IAA17* for cloning into pSAT-nEYFP and pSAT-cEYFP plasmids for subsequent BiFC assay

| | Forward primer | Reverse primer |
|---|---|---|
| *ARF6* | ATCGGAATTCATGAGATTATCTTCAGCT | ATCGAGATCTATGAGATTATCTTCAGCT |
| *ARF8* | ATCGAGATCTATGAAGCTGTCAACATCTGG | ATCGGGTACCCTAGAGATGGGTCGGGTTTTG |
| *IAA6* | ATCGAGATCTATGGCAAAGGA AGGTC | ATCGGGTACCTTAATCTTGCTGGAGACC |
| *IAA9* | ATCGAGATCTATGTCCCCGGAAGAGGAGC | ATCGGGTACCTTAAGCTCTCATCTTCGATTT C |
| *IAA17* | ATCGAGATCTATGATGGGCAGTGTCGAGCTG | ATCGGGTACCTCAAGCTCTGCTCTTGCAC |

**Supplemental Table 5:**

Primers used for amplifying promoters of *IAA6*, *IAA9* and *IAA17*

| **Name** | **Forward primer (LP)** | **Reverse primer (RP)** |
|---|---|---|
| *IAA6* | TTTCTTCCCTCAAATATCTAGC | TTATTTCTTCTCTTTTTTCTTTTTG |
| *IAA9* | CACCAGAGATAGAGAGAAAGGAAGAAAGT | TGGATACTCAAGGTTTCTTTAACTGC |
| *IAA17* | CACCTGTAATCATGTAGGCTGGA | TATTAACCTTTCTTCTTCTTTGGTGT |

### SUPPLEMENTAL REFERENCES

**Alonso, J.M., Stepanova, A.N., Leisse, T.J., Kim, C.J., Chen, H., Shinn, P., Stevenson, D.K., Zimmerman, J., Barajas, P., Cheuk, R., et al.** (2003). Genome-wide insertional mutagenesis of Arabidopsis thaliana. Science **301**:653-657.

**Gutierrez, L., Mongelard, G., Flokova, K., Pacurar, D.I., Novak, O., Staswick, P., Kowalczyk, M., Pacurar, M., Demailly, H., Geiss, G., et al**. (2012). Auxin controls Arabidopsis adventitious root initiation by regulating jasmonic acid homeostasis. Plant Cell **24**:2515-2527.

**Overvoorde, P.J., Okushima, Y., Alonso, J.M., Chan, A., Chang, C., Ecker, J.R., Hughes, B., Liu, A., Onodera, C., Quach, H., et al.** (2005). Functional genomic analysis of the AUXIN/INDOLE-3-ACETIC ACID gene family members in Arabidopsis thaliana. Plant Cell **17**:3282-3300.

**Shi, H., Reiter, R.J., Tan, D.X., and Chan, Z.** (2015). INDOLE-3-ACETIC ACID INDUCIBLE 17 positively modulates natural leaf senescence through melatonin-mediated pathway in Arabidopsis. J. Pineal Res. **58**:26-33.

**Tian, Q., Uhlir, N.J., and Reed, J.W.** (2002). Arabidopsis SHY2/IAA3 inhibits auxin-regulated gene expression. Plant Cell 14:301-319.
